# Adenine Nucleotide Translocase regulates the airway epithelium, mitochondrial metabolism and ciliary function

**DOI:** 10.1101/2020.05.18.101378

**Authors:** Corrine R. Kliment, Jennifer M. K. Nguyen, Mary Jane Kaltreider, YaWen Lu, Steven M. Claypool, Josiah E. Radder, Frank C. Sciurba, Yingze Zhang, Alyssa D. Gregory, Pablo A. Iglesias, Venkataramana K. Sidhaye, Douglas N. Robinson

## Abstract

Airway hydration and ciliary function are critical to airway homeostasis and dysregulated in chronic obstructive lung disease (COPD). COPD is the 4^th^ leading cause of death in the US and is impacted by cigarette smoking with no therapeutic options. We utilized a genetic selection approach in the amoeba *Dictyostelium discoideum* as a comparative discovery tool in lung biology to identify genetic protectors from cigarette smoke (CS). Adenine nucleotide translocase (ANT), a mitochondrial ADP/ATP transporter, was protective against CS in *Dictyostelium* and human bronchial epithelial cells. ANT2 gene expression is reduced in lung tissue from COPD patients and in a mouse smoking model. ANT1 and ANT2 overexpression resulted in enhanced oxidative respiration and ATP flux. In addition to ANT’s presence in the mitochondria, ANT1 and ANT2 reside at the plasma membrane in airway epithelial cells and this localization plays a role in how ANTs regulate airway homeostasis. ANT2 overexpression stimulates airway surface liquid hydration by ATP and maintains ciliary beating after CS exposure, which are key functions of the airway. Our study highlights the potential of ANT modulation in protecting from dysfunctional mitochondrial metabolism, airway hydration, and ciliary motility in COPD.

## Introduction

Central to the function of human airway is the ability to maintain a surface hydration layer that allows the cilia to beat rhythmically to clear mucus, particulates, and infectious organisms from the airway passages. When homeostasis of airway hydration and ciliary function are lost, lung diseases such as chronic obstructive pulmonary disease develop [1]. COPD is a highly prevalent disease worldwide and is comprised of airspace enlargement and airway remodeling leading to significant airflow limitations in patients [2]. Morbidity and mortality due to COPD are rising, and it is currently the 4^th^ leading cause of death in the United States according to the CDC [3], with cigarette smoke being a major inciting factor. Despite significant research, no effective therapies have been identified prevent or reverse COPD development. Mechanistic studies in search of unrealized, essential biology are difficult to conduct in the complex tissue of the human lung. To identify genes that are important for COPD pathogenesis, we have leveraged the social amoeba *Dictyostelium discoideum* as a comparative discovery tool to identify a new pathway in lung biology using a cDNA library with follow-up in primary human airway epithelial cells. We have discovered that the canonical inner mitochondrial membrane protein adenine nucleotide translocase (ANT; paralogs ANT1-4 in humans, with ANT1, 2 and 3 present in the lung) not only regulates mitochondrial metabolism but plays a central function in airway epithelial biology, notably airway hydration, which promotes ciliary function. Mitochondrial dysfunction has been linked to airway remodeling in COPD [1]. As transmembrane ADP/ATP transporters, ANTs are a key connection between mitochondria and the cellular cytoplasm. ANTs’ impact on cellular function appear to be cell-type and ANT-paralog specific. ANTs are known to regulate mitochondrial metabolism and cell fate, including senescence [4] and mitophagy [5]. ANT1 deficiency results in a hypertrophic cardiomyopathy, myopathy, and lactic acidosis [6], while increases in ANT1 protect against cardiac ischemia [7–9] and hepatotoxicity [10]. Reduction in ANT2 protects adipocytes from hypoxia and inflammation [11] while leading to cardiomyocyte developmental abnormalities [12]. The role of ANTs in lung function and COPD pathogenesis are unknown. Most significantly, our results reveal that in a role separate from mitochondria, a population of ANT resides at the plasma membrane of airway epithelial cells where it interacts with the chemiosmotic circuit that controls airway hydration and ciliary beat frequency.

## Results

### Adenine nucleotide translocase identified as a genetic protector from cigarette smoke

Cigarette smoke is a primary insult that leads to the development of COPD and airway dysfunction. To identify potential genetic protectors against cigarette smoke, we utilized the model organism *Dictyostelium* as a tool to identify genes that could then be directly studied in human and mouse models of lung disease (**Fig. 1A**). We challenged cDNA library-transformed *Dictyostelium* cells (an expression cDNA library built from vegetative (growth phase) *Dictyostelium* cells [13, 14]) with a cigarette smoke extract (CSE; prepared in *Dictyostelium* growth media) at the EC40 concentration (the CSE concentration resulting in a 40% decrease in growth rate) (**supplementary Fig. S1A, B**). Over ~3 weeks, we selected for ‘winners’ that could grow in the presence of CSE with growth rates comparable to or exceeding that of untreated wild type cells. We then isolated the plasmids and reintroduced them into fresh *Dictyostelium* cells to confirm the suppression effects. From this, we identified the following genes as protectors against 40% cigarette smoke extract: adenine nucleotide translocase (ANT; encoded by *ancA* in *Dictyostelium*; **Fig. 1B**), the actin-interacting protein-1 (AIP1; encoded by *aip1*), heat shock protein-70 (hsp70; encoded by *mhsp70*), and polyadenylate-binding protein 1A (PABP1; encoded by *pabpc1A*). Because the *ancA* plasmid was the strongest protector, ANT became the focus of this study. The recovered ANT cDNA lacked the initial 46 base pairs encoding for the N-terminal amino acids of the first transmembrane span. Expression of full-length *ancA* gene in *Dictyostelium* also offered protection from 40% CSE (**Fig. 1C**). Adenine nucleotide translocase is an ADP/ATP transporter in the inner mitochondrial membrane. The human genome encodes four paralogs (ANT 1-4), with variable tissue expression [15]. ANT1, ANT2 and ANT3 paralogs are expressed in normal human bronchial epithelial cells with ANT2 being predominant (**supplementary Fig. S1C**). ANTs have 71-89% identity between human isoforms and 62-68% percent identity to AncA in *Dictyostelium* (**supplementary Fig. S1D**). We tested and found that human ANT1 and ANT2 overexpression similarly protected human bronchial epithelial cells (HBEKTs) from cigarette smoke-induced cell death (**Fig. 1D,**adenovirus overexpression in **supplementary Fig. S1E, F**), ranging from 10 to 80% CSE. ANT1 overexpression alone may result in a slight increase in baseline growth rate in HBEKTs. ANT overexpression does not result in changes in expression levels of the mitochondrial proteins, TOM20, VDAC and COX4 (**supplementary Fig. S1E**). We observed no change in nuclei number per cell (single nuclei per cell), indicating no inhibition of cytokinesis. This protection via ANT1 and ANT2 in HBEKT cells occurs in part through a reduction in apoptotic and necrotic cell death in response to CSE (**Fig. 1E, F;** images in **supplementary Fig. S1G**).

**Fig 1.**
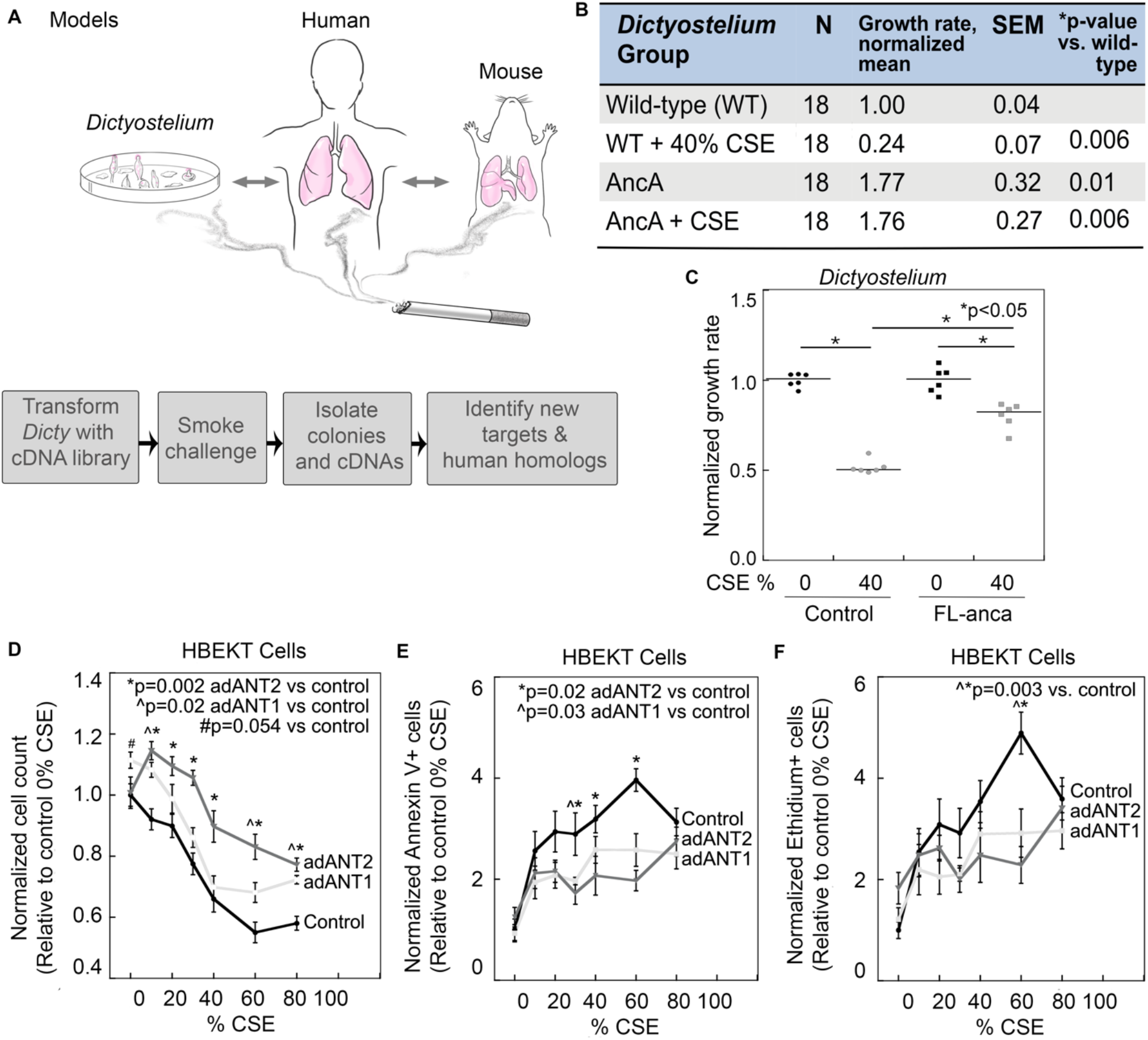
ANT protects against cigarette smoke-induced cell death. **A)** Models and experimental design leveraging *Dictyostelium* to identify relevant pathways in more complex mammalian systems. **B)** *Dictyostelium* cDNA selection growth curve, n=12-18 per group, 6 growth courses. Table includes summative *Dictyostelium* growth rates and statistical parameters. **C)** Growth rates for *Dictyostelium* transformed with full-length *ancA* (FL-*anca*) compared to control plasmid with and without 40% CSE. Median bars are shown. Statistics by ANOVA with Fisher’s LSD post-test. p<0.05 shown on the graph. Cell viability, apoptosis and necrosis were determined in human bronchial epithelial cells (HBEKTs) after 24 hr of 10-80% CSE. **D)** Cell count quantification normalized to 0% CSE, **E**) Apoptosis assessed by annexin-V staining after CSE, **F**) Necrosis assessed by ethidium positive cells after CSE. Data show mean ± SEM, n = 16 from 2-3 experiments. Representative images in **Supplemental Fig. S1**. Statistics by ANOVA; p<0.05 listed on graphs.

### ANT expression level is reduced in COPD

We next determined ANT isoform gene expression in human lung tissue from normal versus COPD subjects in GWAS data from the Lung Genome Research Consortium. *SLC25A4* (ANT1) and *SLC25A5* (ANT2) gene expression in lung was significantly reduced in COPD subjects (**Fig. 2A**). *SLC25A6* (ANT3) was not present in this data set. This is further supported by gene expression analysis by real time PCR of a separate cohort of patients with and without COPD, which revealed a significant reduction in *SLC25A5* (ANT2) and a decreasing trend in *SLC25A4* (ANT1) expression in COPD lung tissue (**Fig. 2B)**. *SLC25A5* (ANT2) gene expression was also significantly reduced in ciliated small airway cells isolated from smokers compared to non-smokers in a publicly available GEO data set, with a decreasing trend for SLC25A4 (**Fig. 2C**). Then, in a mouse model of cigarette smoke exposure for 6 months (when mice develop airspace enlargement and airway remodeling), *slc25a4* (ant1) and *slc25a5* (ant2) gene expression by real time PCR show a downward trend in lungs of C57BL/6 mice compared to air-exposed controls (**Fig. 2D**). Finally, we tested primary normal human bronchial epithelial (NHBE) cells and found that upon smoke exposure, *SLC25A4* (ANT1) trended towards reduced expression but did not reach statistical significance (**Fig. 2E**). These collective observations support the notion that ANTs, and most notably ANT2, are impacted by cigarette smoke exposure and that the reduction in their expression are potential contributors to COPD pathogenesis.

**Fig. 2.**
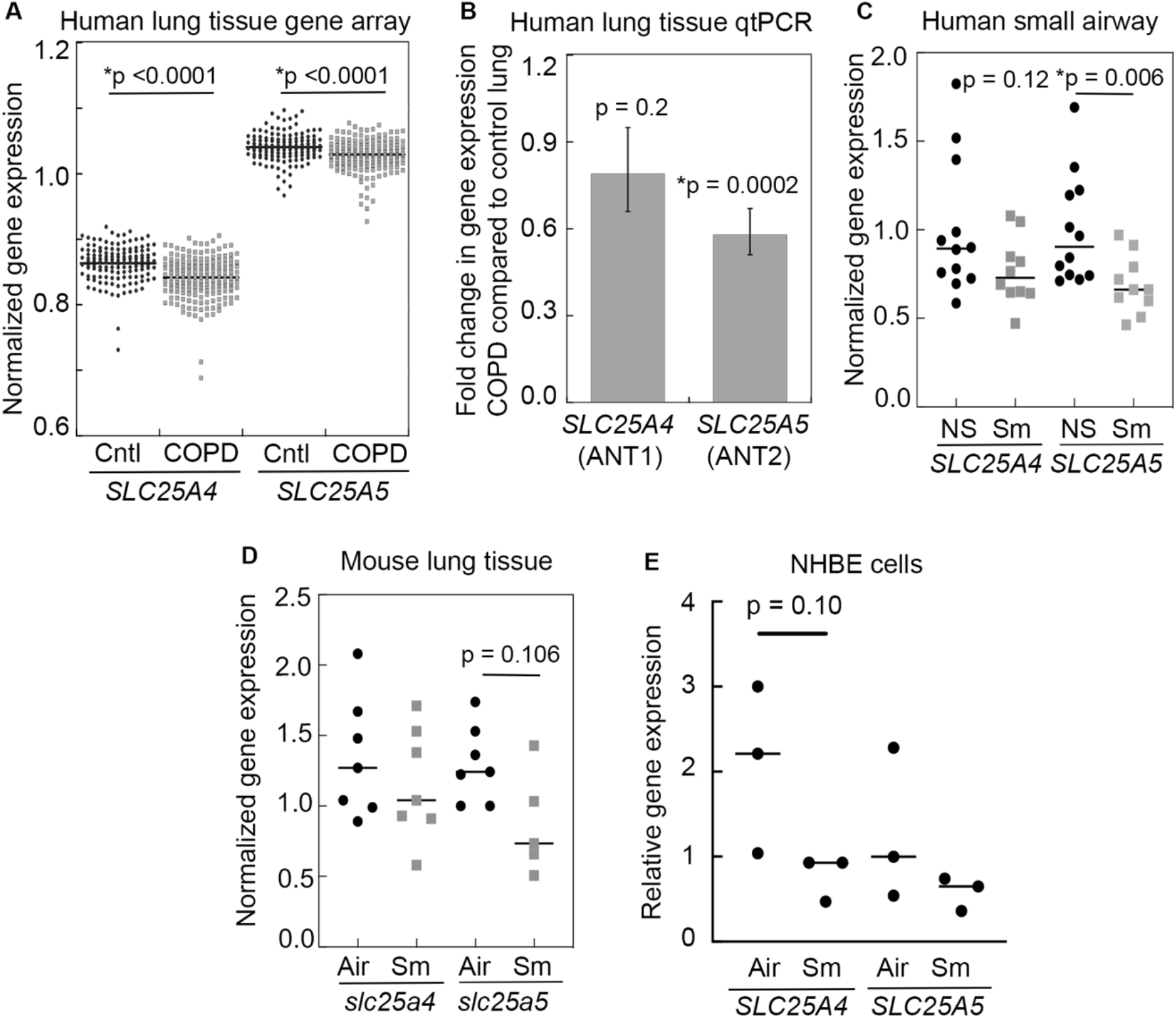
ANT expression is reduced in COPD patients, smokers and a mouse model of COPD. **A)** Human GWAS gene expression data for *SLC25A4* (ANT1) and *SLC25A5* (ANT2) in normal (n=137 subjects) versus COPD (n=219) whole lung tissue, normalized to GPI. Median bars are shown. Statistics by Student’s t-test; p-values noted. **B)** Real-time PCR gene expression for *SLC25A4* and *SLC25A5* in human whole lung tissue from normal (n=35-48) versus COPD (n = 20-23) subjects; Normalized to β-actin and data shown as fold change in COPD compared to normal. Statistics by Student’s t-test; p-values noted. **C)** GWAS gene expression data for *SLC25A4* and *SLC25A5* in human small airway epithelial cells from non-smokers (n=12 subjects) versus smokers (n=10), normalized to GPI, GEO database GDS2486. Medians are shown. Statistics by Mann Whitney; p-values noted. **D)** *slc25a4* (ant1) and *slc25a5* (ant2) gene expression in air versus smoke-treated mouse lungs (6-month exposure), Normalized delta Ct, n=7 per group, normalized to GAPDH. Statistics by Mann Whitney; p-values noted. **E)** Human *SLC25A4* and *SLC25A5* gene expression in air versus Vitrocell smoke-treated NHBE cells (3 exposures with 2 cigarettes per exposure). n = 3 inserts per group. Statistics by Mann Whitney; p-values noted.

### ANT regulates cellular respiration and ATP production

Cigarette smoke is thought to cause an oxidative insult to lung epithelium and mitochondria, resulting in metabolic dysfunction and oxidative stress [1, 16]. ANTs are ADP/ATP transporters that reside in the inner mitochondrial membrane where they provide the source of ADP substrate for the ATP synthase to generate ATP [15]. The ANTs then return the ATP to the cytoplasm where the ATP is utilized as the energy currency of the cell. ANT2 can also reverse this direction to provide ATP substrates to assist the ATP synthase to restore the proton-gradient across the mitochondrial inner membrane. Given the changes in ANT expression found in COPD lung tissue and smoke-exposed lung tissue and cells (**Fig. 2**) and that cigarette smoke can cause mitochondrial dysfunction [1], we tested whether ANT modulation impacts the metabolic activity of intact airway epithelial cells, as well as the impact of CSE on the energy state of the cell. By tracking the oxygen consumption rate (OCR) following sequential addition of inhibitors that target different aspects of the OXPHOS system, we determined the basal OCR (OCR in absence of any inhibitors), mitochondrial ATP production (OCR that is sensitive to complex V inhibition with oligomycin), spare respiratory capacity (difference between maximal OCR in presence of protonophore, FCCP, and basal respiration), and proton leak (difference between OCR upon addition of oligomycin versus OCR when complex I and III inhibitors, rotenone and antimycin A, are added). In addition, the extracellular acidification rate (ECAR) provides insight into the relative importance of OXPHOS versus glycolysis for cellular ATP production (increased production of lactate when glycolytic demand is high).

We found that overexpression at the protein level of ANT1 (~4.3-fold) or ANT2 (to similar expression levels as ANT1) in HBEKT cells (**supplementary Fig. S1E, F**) resulted in increased basal oxygen consumption (26 ± 5.7% and 22 ± 6.1%, respectively) and higher spare respiratory capacity (14 ± 7.2% and 23 ± 6.2%, respectively) in HBEKT cells (**Fig. 3A; also see supplementary Fig. S2A, B**). These increases led to a cellular phenotype characterized by enhanced mitochondrial respiration and glycolytic flux (**Fig. 3B**). Notably, ANT1 overexpression also resulted in enhanced proton leak not seen with ANT2 (**supplementary Fig. S2C**). Overall, CSE had a statistically significant impact on the metabolic state of the cell, decreasing maximal oxygen consumption by 24 ± 2.8% (**supplementary Fig. S2A, B**). The enhanced OCR due to ANT (baseline and maximal OCR) was maintained after 4 hours of CSE exposure compared to an associated drop in respiration in control cells (**supplementary Fig. S2A, B**). The increase in aerobic respiration for ANT2 was in turn reflected in an increase in mitochondrial ATP production (oligomycin-sensitive respiration), 22 ± 5.6% over control (**Fig. 3C**). However, the steady state intracellular ATP concentrations in HBEKT cells with ANT2 overexpression were unchanged at 8 mM (**supplementary Fig. S2G**). Interestingly, this concentration is at the higher end of the range measured for various cell types.

**Fig. 3.**
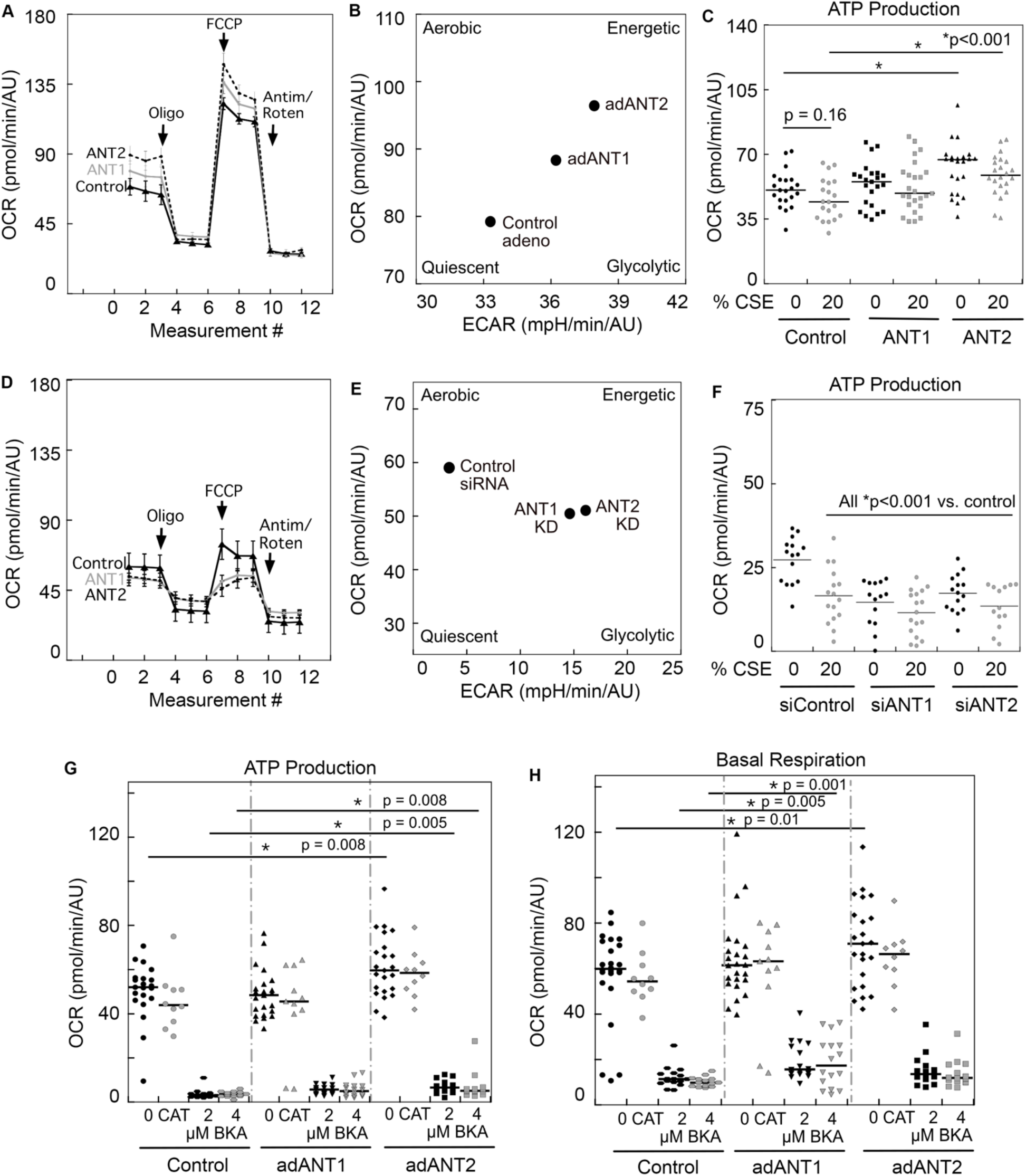
ANT regulates respiration and ATP production in human bronchial epithelial cells. HBEKT cellular metabolism measured using Seahorse Metabolic Analysis. Oxygen consumption rate (OCR) time course with **A)** ANT overexpression or **D)** siRNA suppression. Data show mean ± SEM. OCR versus extracellular acidification rate (ECAR) show **B)** a shift towards a more energetic state with ANT overexpression and **E)** a shift towards a less aerobic state with siRNA suppression. ATP production with **C)** ANT overexpression or **F)** siRNA suppression. Data show median bars (dot plots), n = 15-26 wells from 3 experiments. ANT was overexpressed using adenovirus in HBEKT cells with or without 20 μM CATR or 2 and 4 μM BKA with Seahorse analysis showing **G)** ATP production and **H)** basal OCR. Statistics by ANOVA; *p<0.05 unless noted specifically. Medians are provided in dot plots.

ANT1 and ANT2 siRNA knockdown also resulted in significant reduction in basal oxygen consumption, maximal OCR, and ATP production (**Fig. 3D, F**; **supplementary Fig. S2D, E**), and increased glycolysis (**Fig. 3E**). We observed no significant change in proton leak with ANT1 or ANT2 siRNA knockdown (**supplementary Fig. S2F**). We did observe some effect of siRNA treatment alone on basal OCR levels in these cells. Suppression of ANTs using siRNA for each isoform results in a 61% reduction in ANT1 protein levels and a 74% reduction in ANT2 protein levels (**supplementary Figure S2H**). ANT2 siRNA appears to also partially reduce ANT1 levels (**supplementary Figure S2H**). Protein expression of mitochondrial proteins, TOM20 (translocase of the outer mitochondrial membrane), VDAC (outer mitochondrial membrane) and COX4 (inner mitochondrial membrane) did not change with ANT1 or ANT2 overexpression (**supplementary Fig. S1E**) or suppression (**supplementary Fig. S2J**), suggesting that the metabolic changes seen were not due to changes in these mitochondrial proteins. Other studies have found similar stability with oxidative phosphorylation proteins [17]. Two inhibitors of ANT exist, both blocking the ADP/ATP transport function: carboxyatractyloside (CATR) which is cell membrane-impermeable and bongkrekic acid (BKA), which is membrane-permeable. ATP production and basal oxidative respiration were halted after treatment with the ANT-specific inhibitor BKA, but CATR had no effect (**Fig. 3G, H**), which was anticipated due to CATR’s membrane-impermeability. We noted that BKA inhibited basal respiration more than oligomycin. This could result from the ability of BKA to block ANT-mediated proton leak and/or indicate that ANTs exert greater degree of respiratory control in HBEKTs. Thus, ANTs, predominantly ANT2, appear to protect cells from CSE in part by enhancing both mitochondrial respiration and glycolytic flux of the cells. This in turn leads to improved cell survival in the context of CSE.

Studies suggest that ANT influences the cellular oxidative state [4, 18]. ANT1 deficiency leads to increased cardiac oxidative stress and cardiomyopathy [6, 19], while overexpression is protective against cardiac ischemic injury [7–9]. Overexpression of ANT2 is protective against oxidative stress in hepatocytes [10]. Therefore, we examined whether changes in ANT alters superoxide levels in mitochondria in the context of CSE exposure. In our studies, HBEKT cells did not experience a significant increase in reactive oxygen species (ROS), but ANT1 and 2 expression slightly increased the MitoSOX intensity (mitochondrial ROS reporter) (**supplementary Fig. S2I**). However, it should be noted that this experiment was performed using a high content imager, allowing it to be so heavily powered that a few percent change can be statistically significant. Nevertheless, we suspect that this slight elevation in ROS from ANT1 and ANT2 may reflect increased flux through the electron transport chain resulting in the elevation of ATP production from expression of these proteins.

### ANT2 enhances airway surface hydration and ciliary beat frequency

One of the key functions of the airway epithelium is to clear particulates out of the lung through mucociliary clearance and ciliary beating, which are dysfunctional in COPD due to cigarette smoke [20, 21]. The normal human airway is lined with ciliated airway epithelial and mucosal cells with an overlying airway surface liquid (ASL) layer [22]. The hydration status of the ASL and coordinated ciliary beating are important in maintaining adequate clearance and airway homeostasis. Cilia depend on ATP, not only for energy, but also as a signaling molecule that influences airway surface hydration [23–25]. ASL hydration is important for reduction of mucus viscoelasticity, mucus clearance, and proper ciliary function, which are negatively impacted by cigarette smoke and dysfunctional in COPD [21, 26–29].

We utilized primary ciliated normal human bronchial epithelial cells (NHBEs) and HBEKTs grown at Air-Liquid Interface (ALI) [16] to determine the impact of ANT on ASL thickness and ciliary beat frequency in cells over-expressing ANT1 and ANT2. The primary NHBEs were isolated from human lungs and differentiated to generate cilia with a mucous layer under ALI conditions, recapitulating the human airway. ANT2 overexpression by adenovirus in NHBEs resulted in a 2.3-fold increase in ASL thickness compared to control and ANT1-overexpression cells (**Fig. 4A**). Stable overexpression of ANT2 using lentivirus (**Fig 4B**) similarly yielded an increase in ASL hydration. Importantly, this increase in ASL hydration thickness was inhibited by the cell membrane-impermeable CATR (**Fig. 4C**). With adenoviral overexpression of ANT2, both CATR and BKA (a cell membrane-permeable ANT inhibitor) also inhibited the increase in ASL (**Fig. 4D, E**), which is in contrast to the differential impact of the compounds on metabolic activity where only BKA provided inhibition (**Fig. 3G, H**). In addition, treatment with apyrase to remove extracellular ATP reversed the change in ASL with ANT2 overexpression. These findings suggest that enhanced ASL is mediated through extracellular ATP. Control virus-treated cells treated with CATR also trended towards decreased ASL, suggesting that endogenous ANT may help mediate basal airway ASL homeostasis. ANT2 also enhanced ASL height in HBEKT cells, a non-ciliated bronchial epithelial immortalized cell line, a response which is again abolished by treatment with BKA and CATR (**supplementary Fig. S3A, B**). This observation indicates that ANT2 is important for ASL regulation even in non-differentiated cells.

**Fig. 4.**
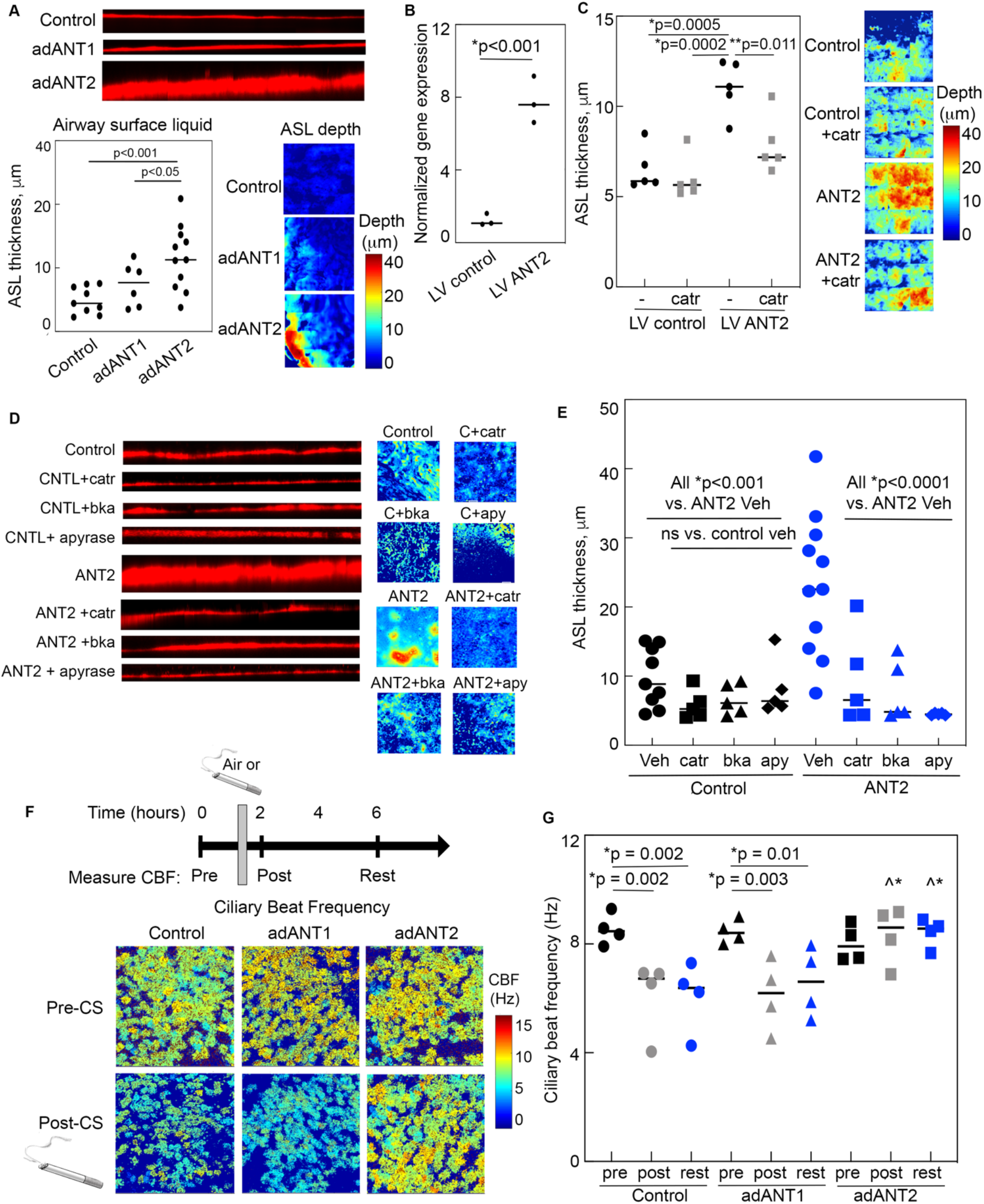
ANT2 enhances ASL hydration and preserves ciliary beat frequency after cigarette smoke. **A)** ASL height in NHBE cells with control adenovirus or overexpression of human ANT1 or ANT2. Representative z-stack of orthogonal views of ASL height with depth heat maps. n = 6-11 ALI cell inserts from 3 different humans. **B)***SLC25A5* (ANT2) gene expression after lentiviral overexpression of ANT2 in primary NHBEs. **C)** ASL height after lentiviral overexpression of ANT2 with or without treatment with 20 μM carboxyatractyloside (CATR). Representative depth heat maps shown. n = 5 ALI inserts. ASL height was determined in primary NHBE cells with adenoviral overexpression of ANT2 versus control virus with PBS vehicle, 20 μM CATR, 4 μM bongkrekic acid (BKA) or apyrase (apy). **D)** Representative ASL orthogonal views and heat maps with **E)** ASL quantification. n = 5-11 inserts from 3 different normal patients over 2-4 days. **F,G)** Quantification of ciliary beat frequency (CBF) measured in control and ANT1 and ANT2 overexpressing NHBEs with pre-treatment (air or CS), 30 min post-treatment, or 4 hr rest. **F)** Experimental setup and example CBF heat maps are shown. CBF from air treated inserts are shown in **Fig. S3**. **G)** Quantification of CBF. P-values determined by ANOVA; *p<0.05, ^*p<0.01. For panels **A**, **B**, **C**, **E**, and **G**, medians are provided.

Because ANT2 overexpression increases and protects ASL height, we hypothesized that ANT2 might also protect ciliated airway epithelial cells from a reduction in ciliary beat frequency (CBF) caused by exposure to acute cigarette smoke. In fact, ANT2 also protected CBF and this protection persisted after a rest period (**Fig. 4F, G**; **supplementary Fig. S4C**). ANT overexpression did not alter baseline CBF, and air exposure alone did not significantly alter CBF in all groups (**supplementary Fig. S3D, E).** Videos of ciliary beating pre- and post-CS exposure for control, ANT1 and ANT2 overexpression cells are provided **(supplementary videos S1-6)**.

### ANT localizes to the plasma membrane in ciliated airway epithelium

Because membrane-impermeable CATR inhibited ANT2’s enhancement of ASL but not its effect on mitochondrial respiration, we hypothesized that a population of ANT likely resides in the plasma membrane where it mediates this ATP-dependent impact on ASL. Literature indicates that ATP levels in the extracellular ASL of airway epithelial cells are ~4-10 nM [23, 24, 30, 31], which when compared to the ~8 mM total intracellular ATP measured for HBEKT cells (**supplementary Fig. S2G**), results in a ~10^5^-10^6^-fold concentration gradient across the plasma membrane. This differential is much steeper than the gradients of nearly all other ions involved in the chemiosmotic cycle. With such a steep gradient, different regulatory mechanisms likely modulate ATP movement through transporters such as the ANTs at the plasma membrane versus what occurs at the mitochondrial membrane.

Therefore, we next examined the cellular distribution of ANTs in the airways of human and mouse lung tissue to determine how ANT expression may vary at baseline and in the context of COPD. As expected, a population of ANT was present near the apical side of the pseudo-stratified columnar cells at the airway lumen, where mitochondria are enriched (**Fig. 5A**, **arrowheads**). This co-localized with TOM20, which is a marker for mitochondria (ANT1 in **Fig. 5B**, **arrowhead**). Unexpectedly, we also observed a population of ANT located outside the boundaries of mitochondria at the region of the apical plasma membrane (**Fig. 5A**, **arrow**). Mitochondrial membrane proteins including the F1-ATPase and ANT have been observed at the plasma membrane in endothelial cells [32, 33] and hepatocytes [34], respectively; however, ANTs localization in lung epithelium was unknown. In human airway epithelial cells, the population of plasma membrane ANT co-localizes with acetylated α-tubulin, which is highly enriched in the cilia (**Fig. 5C**, **arrow**) with a population of ANT that appears to reside at the plasma membrane (**Fig. 5C**, **arrowhead**). We found no evidence of fluorescence bleed through for either α-tubulin or TOM20 (**supplementary Fig. S4A**). Similar colocalization of ANT2/3 with acetylated α-tubulin and TOM20 was also observed (**supplementary Fig. S4B**). We detected this plasma membrane distribution with six different antibodies raised against ANT1 and ANT2 in human airway tissue from normal human lung. The full antibody panel may be found in **supplementary Fig. S5**(with a representative analysis **supplementary Fig. S5B** and ciliary layer vs. mitochondrial quantification **supplementary Fig. S5C, D**). We found differential specificity for the various ANT isoforms by western blotting against yeast expressing each of the human isoforms, ANT1-4 (**supplementary Fig. S6**). To further confirm localization of ANT at the plasma membrane in ciliated cells, we observed populations of native ANT1 and ANT2 that reside in a cellular region similar to nephrocystin-4 (NPHP4), a protein found at the transition zone of cilia (**supplementary Fig. S4C**). ANT staining extends past NPHP4 to the end of the cilia. This suggests that a population of ANTs that reside in an apical cellular region that are separate from mitochondria.

**Fig. 5.**
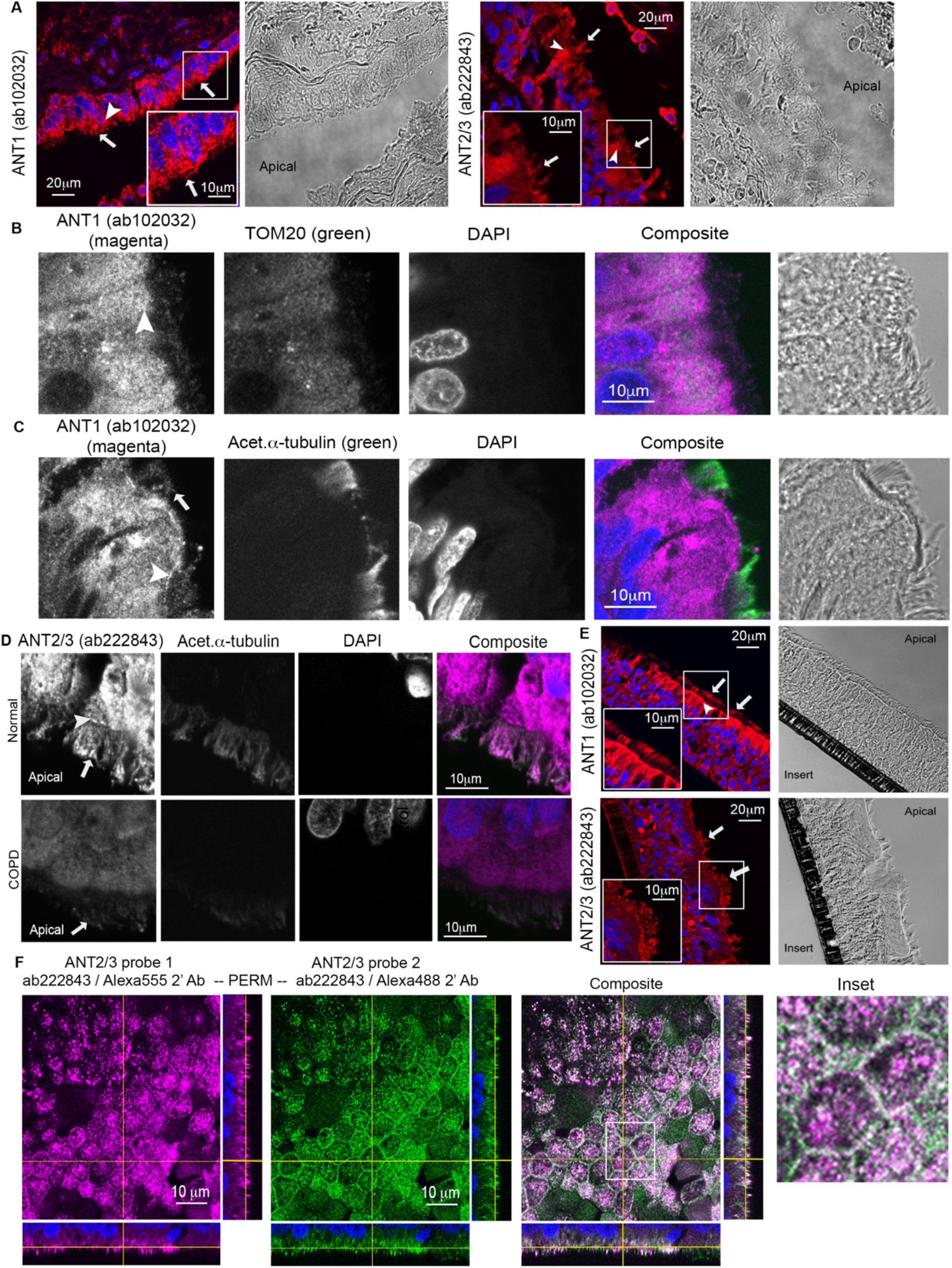
ANT localizes to the plasma membrane in ciliated airway epithelium. **A)** Normal human airway tissue was stained for ANT1 (ab102032) or ANT2/3 (ab222843) and imaged by confocal microscopy. ANT (shown in red) localizes to the mitochondria (arrowhead) and to the ciliary layer (full arrow). The white inset shows additional detail. Representative of n=3-4 different subjects. **B)** ANT1 (magenta, stained in 555 channel) colocalization with TOM20 (green, stained in 647 channel) noted by an arrowhead, imaged by confocal microscopy. Scale bar, 10 μm. **C)** ANT1 (magenta, stained in 555 channel) colocalization with acetylated tubulin (green, stained in 647 channel) at the plasma membrane (arrowhead) and ciliary layer (full arrow), imaged by confocal microscopy. Scale bar, 10 μm. **D)** Human lung tissue from control and COPD patients stained for ANT1 (magenta, 555 channel) and acetylated tubulin (647 channel), imaged by confocal microscopy. Representative of n=3-5 subject per group. **E)** NHBEs grown at air liquid interface stained for native ANT1 or ANT2/3, demonstrating localization of ANT to the mitochondrial layer (arrowhead) and ciliary layer (full arrow). The white inset shows additional detail. **F)** ANT2/3 localizes to the plasma membrane in NHBE cells. NHBE cells were stained for ANT2/3 (without prior permeabilization) followed by secondary antibody Alexa 555 (magenta), then permeabilized and repeat staining with anti-ANT2/3 followed by a secondary antibody Alexa 488 (green). The white is enlarged on the right. Scale bar, 10 μm. n=3 inserts. Images of cells where no permeabilization was used may be found in supplementary Fig. S4E.

In human COPD airway tissue, ANT1 and ANT2 signal is reduced in mitochondrial and ciliary regions compared to normal control tissue (**Fig. 5D,** quantification in **supplementary Fig. S4D**). Similarly, we observed ANTs at the plasma membrane, including in primary ciliated NHBEs grown on air liquid interface derived from four separate human subjects (**Fig. 5E,** panel of antibodies tested in **supplementary Fig. S7**). To confirm the plasma membrane localization of ANTs, ALI cultures were probed first with an antibody to ANT2/3 followed by a secondary Alexa 555 antibody, with or without permeabilization, then stained again with the same primary antibody (secondary Alexa 488). ANT2 was concentrated at the plasma membrane outlining the cell boarders at the apical surface in ALI cultures in both permeabilized and non-permeabilized samples (**Fig. 5F; supplementary Fig. S4E, F)** with similar findings for ANT1 **(supplementary Fig. S4G**). We then drove expression of ANT2-GFP using lentivirus in NHBE cells which were then differentiated at ALI. GFP signal reflecting ANT2-GFP at cilia was again detected and co-localized with acetylated α-tubulin staining (**Fig. 6A**), but when control lentivirus-mediated expression of GFP alone was used, this GFP signal at the cilia was absent.

**Fig. 6.**
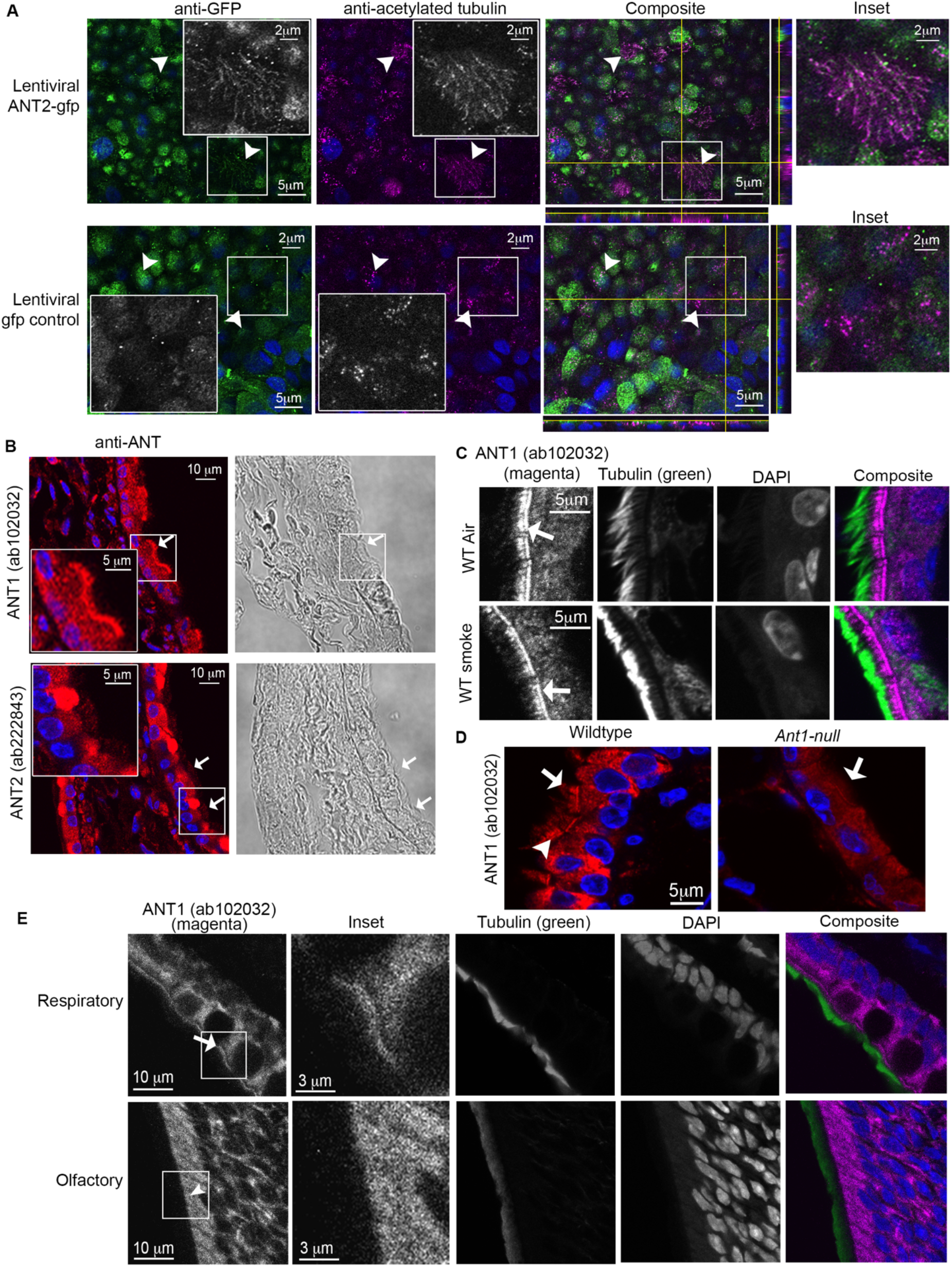
ANT localizes to the plasma membrane in primary human bronchial epithelial cells and mouse airway epithelium. **A)** Lentiviral overexpression of ANT2-GFP and GFP control vector in differentiated NHBE cells. Cells were stained with anti-GFP (green, Alexa 488) and anti-acetylated tubulin (magenta, Alexa 647) and imaged with confocal microscopy. Ciliated cells are noted with an arrowhead. Scale bar, 5 μm. Grayscale insets are included for each color channel. The composite color inset is enlarged on the right. **B)** Normal mouse airway tissue was stained for ANT1 (ab102032) or ANT2/3 (ab222843) and imaged by confocal microscopy. ANT (shown in red) localizes to the mitochondria and to the plasma membrane/ciliary layer (full arrow). The white inset shows additional detail, scale bar 5 μm. Representative of n=3. **C)** Mouse lungs (air versus smoke-exposed for 6 months) stained for ANT1 (magenta, ab102032), acetylated α-tubulin (green) and DAPI (blue). n=5 mice per group. Scale bar, 5 μm. **D)** Staining of wildtype C57BL/6EiJ and *ant1* null mouse airways for ANT1 (ab102032). Plasma membrane localization is noted by the arrowhead and ciliary layers by the full arrow. Scale bar, 5 μm. **E)** ANT1 (magenta) and acetylated α-tubulin (green) in mouse upper respiratory and olfactory epithelium, scale bar 10 μm. Full arrows note apical plasma membrane ANT while arrowheads note mitochondrial ANT. Apical membrane ANT is present in motile ciliated respiratory epithelium compared to non-motile ciliated olfactory epithelium. The white inset is enlarged in the panel, scale bar 3 μm.

Mice have ANT paralogs 1, 2 and 4 with similar cell type expression to humans. We looked at ant1 and ant2 localization in the mouse airway epithelium and found murine ANT present at the plasma membrane of airway cells (**Fig. 6B**). Structured illumination microscopy of ant1 in mouse airway epithelium indicated a highly patterned distribution along apical cell surface, the base of cilia and the periciliary membrane (**Fig. 6C; supplementary video S7**). In mice exposed to cigarette smoke for 6 months, this ant pattern was disrupted with less organized ant distribution at the plasma membrane compared to air-treated control mice (**Fig. 6C; supplementary video S8**). Further, airway epithelial cells from *ant1*-null mice [6] had significantly less plasma membrane staining compared to wildtype mice using ant1 antibody ab102032 (**Fig. 6D**). However, the remaining staining existed in a similar pattern, suggesting that this antibody also partially recognizes the ant2 paralog. The plasma membrane localization of ant1 was not a generalized pattern in all ciliated cells. Non-motile cilia of the olfactory epithelium in mice did not have plasma membrane ant while neighboring motile respiratory epithelia contain ant at the apical plasma membrane in the same mouse tissue sections (**Fig. 6E**).

In summary, this discovery of a population of ANT at the plasma membrane and the effect of ANT2 on ASL and CBF suggests that ANT may also regulate ATP at the plasma membrane. Pannexin has already been implicated in providing part of the ATP transport in normal airways [31, 35, 36]. The population of ANT at the plasma membrane, which CATR can inhibit, may provide another mechanism for ATP transport at the airway surface, which in turn promotes an increase in ASL height and protection of ciliary beating when challenged by cigarette smoke.

## Discussion

By utilizing a powerful model organism platform extending from *Dictyostelium discoideum* to human lung disease, we have discovered that the canonical inner mitochondrial membrane protein adenine nucleotide translocase (ANT) plays a central function in airway epithelial biology. Namely, ANT serves multiple roles including protection of cell viability by enhancing mitochondrial respiration, promotion of airway hydration, and preservation of ciliary function (**Fig. 7**). Prior studies suggest an important role for mitochondrial dysfunction due to cigarette smoke and COPD [1, 16]. Here, we find that by manipulating ANT expression, we can increase mitochondrial respiration and glycolytic flux, which allows cells to withstand injury and prevent subsequent cell death in the setting of insults such as cigarette smoke. ANT’s metabolic role may be primarily responsible for this protection because ANT can protect the growth of *Dictyostelium* cells, which do not have cilia and no apparent plasma membrane-associated population of ANT. Expression levels of ANT appear to help guide overall cellular respiration and ATP production in airway epithelial cells. In further support of the key role of ANT2 in airway biology and COPD, we find that *SLC25A5* (ANT2) gene expression is reduced in the lungs of COPD subjects from separate human tissue cohorts, specifically in human ciliated airway epithelial cells from smokers, and a mouse model of smoke exposure. This highlights the importance of understanding how ANTs impact COPD pathogenesis.

**Fig. 7.**
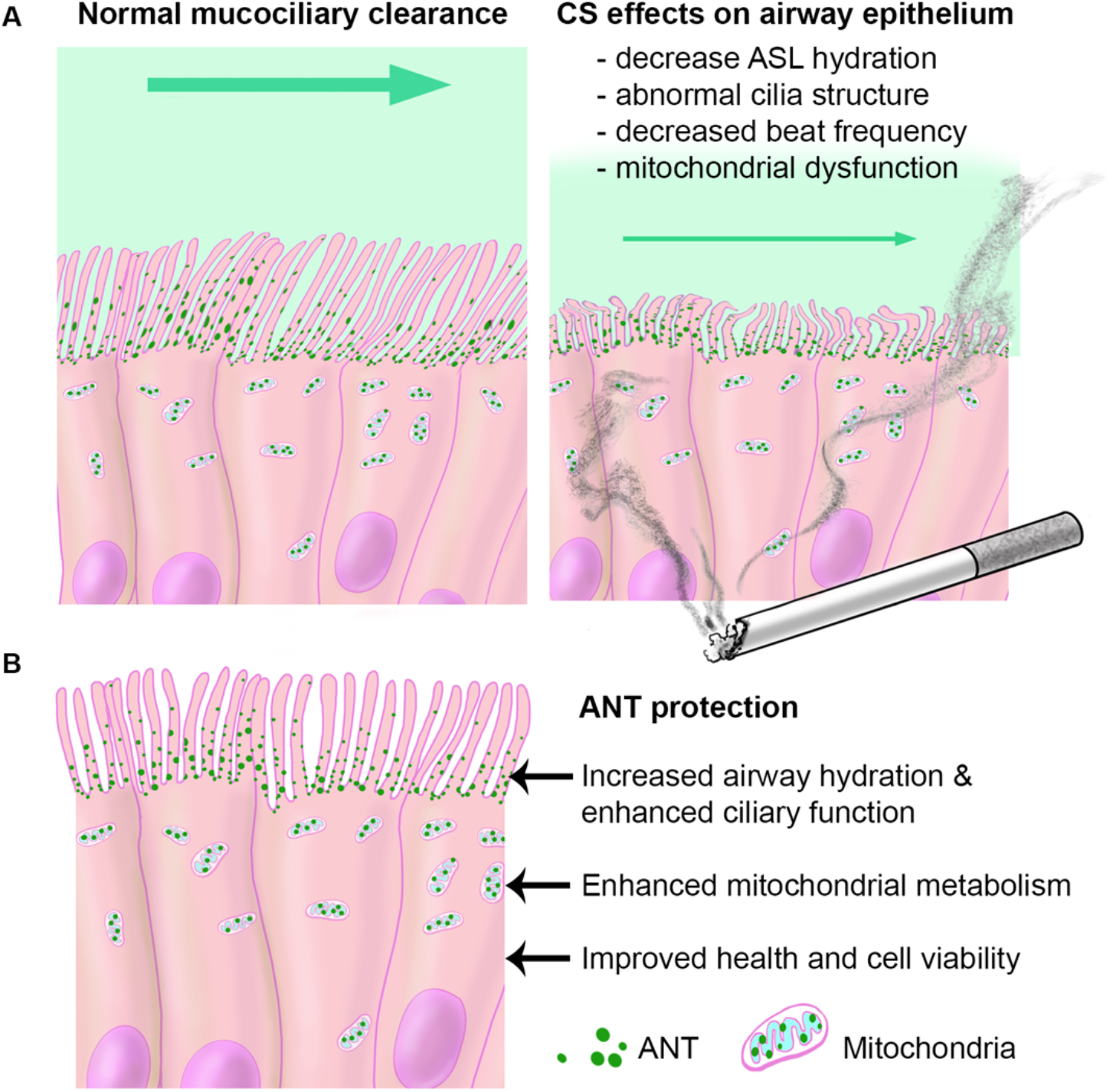
ANT localizes to motile cilia where ANT2 is utilized to regulate airway surface hydration and preserve ciliary function. **A)** Normal mucociliary transport requires a delicate balance of airway surface hydration and ciliary beating to move mucus and particulates out of the lungs. Cigarette smoke leads to decreased ASL [26], decreased ciliary beat frequency (current data and [29]) and abnormal cilia structure [21, 27]; which contribute to reduced mucociliary clearance and development of lung disease. **B)** ANT2 localizes to the plasma membrane in ciliated airway epithelial cells. It protects human airway epithelium from CS-induced injury by increasing airway surface hydration through ATP regulation, preserving ciliary beat frequency, enhancing metabolism and maintaining cell viability.

Classically, ANTs are translated in the cytoplasm and imported to the inner mitochondrial membrane where they transport ADP and ATP. However, we found that populations of ANT1 and ANT2 are also present in the plasma membrane of the airway epithelium. These observations are based upon three major lines of evidence: 1. Inhibition of ASL thickness modulation with a cell-impermeable ANT inhibitor CATR; 2. Immunocytochemistry and immunohistochemistry with a library of anti-ANT1 and anti-ANT2 antibodies; and 3. Expression of ANT-GFP fusion proteins in primary airway epithelial cells. These observations support the conclusion that a population of ANT1 and ANT2 resides at the plasma membrane in epithelial cells where they help modulate ASL and ciliary function. Interestingly, both ANT1 and ANT2 contain predicted secretion signal sequences, which could be the basis for their plasma membrane insertion.

One prior study using a proteomics approach identified ANT1 as a potential interactor with UBXD8 (FAF2), a ciliary protein [37]. Our discovery of a subset of ANT that resides at the plasma membrane of airway cells indicates that ANT has important non-mitochondrial functions in primary airway epithelium. ANT2 in particular has a protective role in airway hydration and ciliary beating, two key components of mucociliary clearance that are abnormal when COPD develops in the lung. Mucociliary clearance [38] is critical for removal of toxins, inhaled particulates, and bacteria from the airway, and maintenance of mucus homeostasis is the first line of defense in the lower respiratory tract. The efficiency of mucociliary clearance depends on the hydration status of the airway surface liquid, the ciliary beat frequency, and having appropriate mucus production. Insults such as chronic cigarette smoke exposure lead to decreased ASL levels, reduced CBF, and abnormal cilia [21, 26, 27, 29] (**Fig. 7A**). These functions are disrupted in airway diseases from COPD to cystic fibrosis. We propose that ANT2 resides at the plasma membrane to regulate airway hydration by transport of ATP to the extracellular surface, influencing ciliary beating.

Several physiologic pathways likely exist to regulate ATP levels at the apical surface of the airway epithelium such as pannexin [31]. We propose that ANT acts in concert to maintain ASL height. While we did not see an overwhelming effect of ANT inhibitors on ASL height at baseline, ANT may rather be part of a stress response to insults such as cigarette smoke. The electrogenic exchange of ADP/ATP via ANT is influenced by the proton-motive force across the inner mitochondria membrane. In the context of the plasma membrane, it is likely that ANT transport will be under alternate control as ATP export is likely driven by the steep concentration gradient across the plasma membrane. We propose that the consumption of extracellular ATP then provides the substrate (ADP) needed for ANT to transition back into a conformation that makes it available to transport cytoplasmic ATP, although this awaits formal testing. Future investigations could additionally delineate how ANT inhibitors (specifically CATR) affect other known regulators of ASL hydration and ATP flux such as pannexin, purinergic receptors and cystic fibrosis transmembrane conductance regulator (CFTR).

In summary, ANT2 provides a powerful strategy for altering mitochondrial function by increasing ATP production and manipulating ciliary function through both airway surface hydration and preservation of ciliary beating (**Fig. 7B**). This dynamic interplay between hydration of ciliated epithelial surfaces and coordinated ciliary beating is important for efficient clearance of mucus and particulates out of the lungs. The discovery of two populations of ANT in the lung, mitochondrial and plasma membrane, provide distinct paths that will benefit from additional investigation and could be modulated to alter disease development in COPD. The roles of ANT, most prominently ANT2, in cellular metabolism, epithelial surface hydration and ciliary function will likely have broad applicability and impact in a variety of other lung diseases.

## Materials and Methods

Further information and requests for resources and reagents should be directed to and will be fulfilled by the Lead Contacts, Corrine Kliment and Douglas Robinson.

### EXPERIMENTAL METHODS

#### Genetic selection studies in *Dictyostelium*

Wild type *Dictyostelium discoideum* cells used in our studies were wild-type cells, the HS1000 (Ax3(Rep ORF+7-3)) strain, grown in Hans’ enriched HL-5 media (1.4x HL-5, containing 8% FM (ForMedium, Norfolk, UK) plus 60 U/ml penicillin, 60 μg/ml streptomycin sulfate) and selected for plasmid transformations with 15 μg/ml G418. Cells were propagated at 22°C on 10-cm Petri dish plates. For suspension growth, cells were cultured in 10-ml culture volumes in 125-ml Erlenmeyer flasks at 180 rpm, 22°C. Cell densities were determined by counting cells on a hemocytometer. The expression cDNA library was constructed in the pLD1A15SN plasmid with a G418 resistance cassette and using cDNA generated from mRNA isolated from vegetative (growth phase) *Dictyostelium* cells [13, 14]. For a negative control, a GFP-pLD1A15SN plasmid was used. Transformation of wild-type cells was performed using electroporation using a Gene Pulser (BioRad) and a total of 35,000 clones (35 pools of ~1000 clones/pool) were generated. Transformants were grown in HL-5 supplemented media with G418 alone or with 40% cigarette smoke extract over 3-4 growth cycles. Cells were pulsed one time in a 0.4-cm cuvette at 3 μF. After electroporation, the cells were placed in cold Han’s enriched HL-5 to allow recovery and grown for 24 hr at 22°C followed by replacement of media with Han’s enriched HL-5 with G418. Media was changed every 2-3 days until clones were harvested. For transformation of wild type cells, a total of 35,000 clones (35 pools of ~1000 clones/pool) were generated and subjected to growth selection. Cells were initially cultured at 1×10^5^ cells per mL in HL-5 supplemented media with G418 alone or with 40% cigarette smoke extract over 3-4 growth cycles. Relative growth rates were determined by counting cells using a hemocytometer. Cell densities were plotted versus time to generate a log phase curve that was exponentially fit using KaleidaGraph (Synergy Software). The concentration of CSE that results in a 40% decrease in growth rate is reported as the EC40 concentration. DNA was isolated from cultures demonstrating a growth advantage in the context of cigarette smoke, using a glass milk protocol. DNA was transformed into STBL2 cells and individual clones were selected for DNA clean-up and restriction endonuclease digestion. Recovered DNA was sequenced using standard procedures. cDNA from recovered clones (those showing suppression of the CSE growth phenotype) were transformed into the parent wild-type cells to confirm that the DNA construct could recapitulate the suppression phenotype. Molecular cloning was completed to yield a full length *AncA* plasmid to confirm that suppression of the growth phenotype in the context of 40% CSE in suspension cultures. Primers for full length *AncA* used were included at 1 μM: forward AAAAAAGTCGACATGTCTAACCAAAAGAAAAACGACGTATCTTCATTTG; reverse AAAAAAGCGGCCGCTTATTCAGAACCAACACCACC.

#### Human cell culture (cell line and primary cells)

The human bronchial epithelial cell line (HBEKT, immortalized by Cdk4 and hTERT [39] (a gift from John Minna) was used for cell viability, metabolic analysis, and airway surface liquid studies. Cells were verified to be mycoplasma-negative and genetically authenticated by STR profiling (Gene Print 10, Promega). HBEKT cells were maintained in keratinocyte serum free media with supplementation according to Lonza protocol. Cells were split when 80-90% confluent after trypsinization with 0.05% trypsin and neutralization with trypsin neutralizing solution.

Sources of primary normal human bronchial epithelial cells (NHBEs) include the University of Pittsburgh Airway Cell Core, Lonza and MatTek. NHBEs were used from a total of five different normal human subjects. Cells attained from the University of Pittsburgh cell core were attained as previously described [40, 41] with a protocol approved by the University of Pittsburgh Investigational Review Board. NHBEs were grown to 80-90% confluence in collagen-coated flasks then seeded onto Type I collagen-coated (50 μg/mL in 0.02N acetic acid) transparent PET transwell inserts (0.4-μm pore, 24-well insert size 0.33 cm^2^ and 12-well insert size 1.12 cm^2^, Corning Costar) at a density of ~5-6 × 10^5^ cells/cm^2^. Once a confluent monolayer is formed on the inserts, the apical media is removed and cells are grown at air-liquid interface [16] over three to six weeks for differentiation into ciliated airway epithelium.

NHBE cells from Lonza and MatTek were initially grown in growth media including BEGM media (Lonza) with recommended supplements (bovine pituitary extract, insulin, hydrocortisone, epinephrine, transferrin, recombinant human epidermal growth factor, retinoic acid, triiodothyronine (T3), and gentamicin sulfate amphotericin-B) with additional bovine pituitary extract (12.6 μg/mL, AthenaES), bovine serum albumin (BSA) (final concentration of 1.5 μg/mL, Sigma-Aldrich), retinoic acid (final concentration of 0.1 μM) and epidermal growth factor (final concentration of 25 ng/mL). When NHBE cells were grown at air liquid interface, basal air liquid interface media was used including BEGM media (Lonza) and DMEM with recommended Lonza supplements (1 supplement pack per 500 mL of media) with additional bovine pituitary extract (12.6 μg/mL, Lonza and AthenaES), BSA (final concentration of 1.5 μg/mL, Sigma-Aldrich), and retinoic acid (final concentration of 0.1 μM).

#### *In vitro* cigarette smoke exposure

Methods used for cigarette smoke exposure include prepared cigarette smoke extract (CSE) or gaseous smoke using a Vitrocell exposure chamber. CSE was made using a peristaltic pump that smoked one research grade cigarette (Tobacco Health Research Institute, University of Kentucky, Lexington, KY) over 6 min and bubbled into 25 mL of cell specific media, yielding what was considered 100% CSE, which was then filtered with a 0.22-μm filter. Doses of CSE from 10-80% were made and used within 6 hr. The Vitrocell smoke exposure chamber was used to expose NHBE cells at ALI to humidified air or cigarette smoke. A single exposure is considered air for 16 min or two cigarettes over 16 min using an International Organization for Standardization (ISO) protocol.

#### Targeted gene delivery or suppression

Adenovirus constructs were developed for gene delivery of control eGFP, ANT1-GFP (Ad-h-*SLC25A4*/eGFP, GenBank BC008664.1), and ANT2-GFP (Ad-h-*SLC25A5*/eGFP, GenBank BC056160.1) from Vector Biolabs (Malvern, PA) in NHBE and HBEKT cells. In select ASL studies, lentiviral constructs (Vector Builder Inc) were used in NHBE cells including control GFP (LVM (VB160109-10005) and ANT2-gfp (Cat#: LVS(VB190613-1074dha)-C) at an MOI of 20. NHBE cells were infected with lentivirus for 24 hr at the time of seeding onto transwell inserts, followed by puromycin antibiotic resistance selection for 48 hr. The cells were then differentiated at ALI as described. For each cell type, a multiplicity of infection (MOI) titration was completed. Protein expression and localization were confirmed. Adenoviral and lentiviral infection efficiencies were approximately 85-95% with confirmed expression of ANT-GFP isoforms. Adenoviral infected cells were used for experiments 48 hr after initial virus exposure. siRNA ON-TARGETplus smart pools were used for genetic suppression in HBEKT cells (Dharmacon: ANT1 (*SLC25A4*, #L-007485-00-0005), ANT2 (*SLC25A5*, # L-007486-02-0005), non-targeting control pool #D-001810-10-05) using Lipofectamine 2000 (Invitrogen) per manufacturer’s instructions. Overexpression and targeted knockdown were confirmed with western analysis for each protein at 24, 48 and 72 hr.

#### Cellular Viability, Metabolism and ATP measurements

For viability analysis, HBEKT cells were cultured in 384-well glass bottom plates, and infected with adenovirus constructs for ANT1-GFP, ANT2-GFP and control GFP vector. At 48 hr post-infection, cells were treated with KSFM media alone or 10-80% CSE for 24 hr. Cells were stained with Alexa 405-Annexin V, ethidium homodimer (necrotic cells) (Biotium, Inc), and Draq5 (nuclei) for viability experiments or MitoSOX Red and Draq5 for oxidative stress analysis. Cells were washed once with binding buffer and imaged in L-15 Leibovitz media on a Molecular Devices High Content imager using 20x and 60x objectives capturing 4 frames per well. Mitochondrial reactive oxygen species production was assessed using MitoSOX Red staining in cells treated with adenovirus and CSE as above. Cells were washed once with PBS and incubated with MitoSOX for 10 min at 37°C, 5% CO2. Nuclei were labeled with Draq5 (#4084, Cell Signaling). Cells were washed three times with PBS and placed in L15 media for imaging using the High Content Imager. For cell analysis, fluorescence intensity of mitochondria was assessed with exclusion of the nucleus. Five hundred to seventeen hundred cells were analyzed per well, 8 wells per experiment on 2-3 different days. For cell analysis, MetaXpress software (Molecular Devices) was used to separate cells infected with adenovirus constructs from uninfected cells. The average fluorescence intensities of MitoSOX staining in mitochondria were determined with exclusion of the nucleus. About 300-400 cells were analyzed per well for a total of 700-1500 cells per group. Across all replicates, 2800-4500 cells per group were evaluated.

The Seahorse Mitostress assay (XF96 FluxAnalyzer, Agilent) was used for metabolic analysis of HBEKT cells with ANT modulation. HBEKT cells were cultured in 96-well Seahorse assay plates (Agilent) and infected with adenovirus constructs for ANT1, ANT2 and control for target overexpression at 48 hr (MOI 40, Vector BioLabs) or siRNA target suppression using siRNA and Lipofectamine as described above for ANT1, ANT2 or non-targeting control siRNA at 72 hr. After viral infection or siRNA transfection, cells were treated for 4 hr with KSFM media alone or 20% CSE. Cells were washed once in KSFM, followed by washes in buffered Seahorse Assay medium (pH 7.4) and subsequent metabolic testing according to the manufacturer’s protocol (Agilent Seahorse XF96). Oligomycin (2 μM, ATPase inhibitor), FCCP (0.25 μM mitochondrial uncoupler), and a cocktail of rotenone (0.5 μM, ETC complex I inhibitor) and antimycin A (0.5 μM, ETC complex III inhibitor) were sequentially injected after three basal rates were measured. Real-time oxygen consumption rate (OCR) is determined including basal OCR (before test kit inhibitors), maximal OCR (after FCCP treatment, a mitochondrial uncoupler), ATP production (OCR after injection of the complex V inhibitor oligomycin), and proton leak. Sample measurements were normalized to total cellular mass determined by CyQuant Assay according to the manufacturer protocol (Molecular Probes, ThermoFisher). For experiments including ANT-specific inhibitors, carboxyatractyloside (CATR, 20 μM, Sigma), bongkrekic acid (BKA, 4 μM, Sigma), or PBS vehicle control were injected into wells after basal OCR measurements were taken, followed by repeat OCR measurements 30 min after compound injection. Cells were then assayed according to the Seahorse protocol as described above.

Steady state intracellular ATP concentrations were measured in HBEKT cells after overexpression of adenoviral ANT1-GFP, ANT2-GFP, and GFP control for 48 hr followed by treatment with media alone or 20% CSE for 4 hr. Cell lysates were collected with a ATP lysis buffer with 300 μM of ecto-ATPase inhibitor ARL 67156 (Sigma) using a luciferin-luciferase bioluminescence ATP Determination Kit (ThermoFisher) with an ATP standard curve. The ATP concentration in mM/HBEKT cell was calculated by considering the cellular volume of a single HBEKT cell and the amount of ATP in a single cell as shown in the following equation:

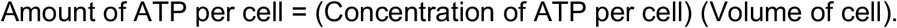

Spherical cell volume was determined after measuring the diameter of trypsinized HBEKT cells on an epifluorescence microscope (Olympus). The amount of ATP for one cell was derived from the amount of protein in one cell found via Bradford assay (BioRad).

#### Measuring Airway Surface Liquid (ASL) Height

ASL was visualized by adding 10 kD-Texas Red-dextran (ThermoFisher) to the apical surface of NHBE or HBEKT cultures at air liquid interface, as previously described [23, 25, 42] and z-stack ASL images collected 4 hr later on confocal microscope at 37°C, 5% CO_2_. The apical surface of cells was washed once with PBS 18 hr prior to the addition of Texas Red. ASL was visualized by adding 10 kD Texas Red-dextran (17.5 μl per 12-well insert and 6.7 μl per 24-well insert at 1.5 mg/mL in sterile PBS, ThermoFisher) to the apical surface of NHBE or HBEKT cultures at air liquid interface, as previously described [23, 25, 42]. Fluorinert (100 μl per 12-well insert and 30 μl per 24-well insert) was added to the top of the insert cultures to prevent evaporation. ASL was imaged 4 hr later by acquiring 3×3 or 4×4 tiled z-stacks by live-cell confocal microscopy with a heat and CO_2_ controlled stage (Zeiss 780 or Nikon A1R confocal microscope with a 40x water objective). Z-stack images at the ASL were captured at a step size of 0.46 μm and were analyzed by segmentation and pixel thickness analysis. For experiments with ANT inhibitors, CATR (20 μM, Sigma) and BKA (4 μM, Sigma) in PBS (or control PBS alone) were added to the Texas Red dextran dye and applied to the apical surface of ALI cultures (NHBEs or HBEKTs) 4 hr prior to ASL assessment. For apyrase treatment, 10 units of apyrase (Sigma) was added with Texas Red-dextran as described above 4 hr prior to ASL assessment.

Thickness across the sample was computed using a custom script, written in Matlab (Mathworks, Natick, MA). Briefly, for each value *x* in the three-dimensional images, Im(*x, y, z*), the resultant (*y, z*) slice was segmented using adaptive thresholding (Matlab command: imbinarize), followed by operations to fill holes (imfill), morphological opening (imopen) and filtering of small regions (bwareaopen) (**supplementary Fig. S8A**). The threshold level was adjusted after preprocessing the complete image and the process repeated. For each y in this binary image, the number of segmented pixels was counted, giving a measure of the thickness in the (x, y) location; see Supplementary Information 1. This gave a histogram of depths over the image (**supplementary Fig. S8B, C**). An average depth was computed for all pixels in which a non-zero depth was detected (to avoid edge effects). The process was repeated by fixing y and working with the (x, z) slice. The differences between the averages were typically less than 1–2%. Thickness in pixels was converted to μm using a slice thickness of 0.46 μm, the step size used to collect the z-stack.

#### Ciliary Beat Frequency

Ciliary beat frequency (CBF) was assessed at 48 hr post-adenoviral infection. Inserts were imaged in a 12-well Falcon plate with Leibovitz L-15 buffered media (Gibco) and imaged within 2 min of placement. Bright field images of beating cilia were captured at 160 frames per sec over 4 seconds (3-5 videos per insert, Leica Spinning disc confocal with 40x water objective). For air or cigarette smoke experiments, “pre-treatment” images (n = 3 per insert) were captured at random insert locations followed by exposure to air or cigarette smoke by the Vitrocell system (2 cigarettes over 16 min). “Post-treatment” images were collected at similar locations on each insert 30 min after the treatment. Cells were kept in ALI growth media at 37°C and 5% CO2 when not being imaged. Cells were “rested” for 4 hr and imaged again (n= 3 videos per insert). Beat frequency analysis was completed using a Matlab script assessing pixel intensity fluctuations (**supplementary Fig. S9**). CBF data was compared with SAVA analysis [43] and both outputs agreed closely.

A custom script, written in Matlab, was used to estimate the beating frequency of cilia. Individual images from a video consisting of *N* frames were used to create a three-dimensional matrix Im_*k*_ (*x, y*), where (*x, y*) denotes location in the image of each pixel, and *k* ∈ 1*, …, n* is the frame number. For each (*x, y*), the corresponding sequence of intensities was first normalized:

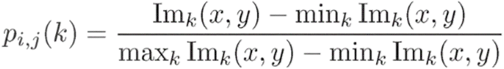

so that 0 ≤ *p_i,j_* (*k*) ≤ 1 and filtered by removing the mean value:

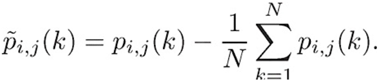

The Fast-Fourier transform (FFT) of 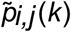 was obtained using the MATLAB command fft:

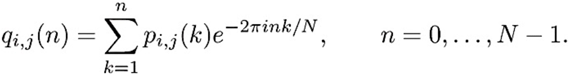

and the corresponding single-sided power spectrum was computed (where *N* is even):

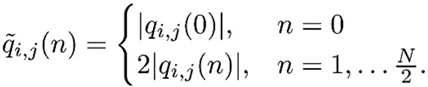

This gives a power spectrum for each pixel (see **supplementary Fig. S9B, C)**.

To determine the frequency of beating, we carried out two approaches. In the first, we found the frequency with highest power density for each of the pixels. We then used a threshold (set at 0.125 A.U.) to determine whether there was any detectable power in that pixel or not. The frequency with highest power was determined, and this data aggregated over all pixels meeting this threshold (**supplementary Fig. S9C**). The data between 2 and 20 Hz of the corresponding histogram was then normalized and fit by a single Gaussian, 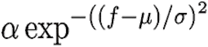 where *f* is the frequency, using the command fit. The value of μ was used as a measure of beating frequency.

In the second method, we aggregated the power spectra from all pixels (with no thresholding): 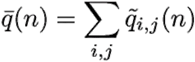 and fit the data between 2 and 20 Hz to a Gaussian mixture model using three modes (**supplementary Fig. S9D**). This resultant μ value with the greatest contribution to the mixture model was used. Both methods provided similar estimates, so the data reported used Method 1 frequencies.

#### Animal studies

##### Mouse smoke exposure and processing

All animal experiments were performed in accordance with the Institutional Animal Care and Use Committee (IACUC) of the University of Pittsburgh. Animals were housed according to standard housing criteria. C57Bl/6J mice (female mice at 10 weeks of age, n = 5 per group) were subjected to the smoke of 4 unfiltered cigarettes per day (lot# 1R5F; University of Kentucky, Lexington, KY), 5 days a week for a duration of 6 months, using a smoking apparatus that delivers targeted cigarette smoke to single mice isolated in individual chambers [44]. The controls in each group were exposed to room-air alone. These mice were caged separately and housed in the same facility as their smoke-exposed counterparts. At the completion of each experiment, mice were killed by CO_2_ inhalation, the chest was opened, and the trachea was cannulated. Lungs were inflated with 10% buffered formalin at a constant pressure of 25 cm H_2_O for 15 min. The lungs were then ligated, excised, and fixed in formalin for 24 hr before embedding in paraffin. Serial midsagittal sections were obtained for histological analysis. In a subset of animals, lungs did not undergo fixation and instead were excised and placed directly in liquid nitrogen for RNA isolation. Whole lung RNA was isolated using Trizol per the manufacturer’s protocol.

##### Human tissue studies

Studies using human tissues were approved by the University of Pittsburgh and Johns Hopkins Institutional Review Boards (IRB) in accordance with ethical guidelines. For immunohistochemical studies, donor human lung tissue (control and COPD patients) was attained and approved through the National Heart, Lung, and Blood Institute’s Lung Tissue Research Consortium database and the Center for Organ Recovery and Education (CORE) at the University of Pittsburgh [40, 41]. Human lung tissues used for real time PCR were obtained from explanted lungs after lung transplantation or donor lungs not suitable for organ transplantation (The Airway Cell and Tissue Core, supported by P30 DK072506, NIDDK and the CFF RDP to the University of Pittsburgh). Lung tissues were stored at −80°C until future usage.

#### Human and Mouse Lung Tissue mRNA Expression

Human lung tissue from The Airway Cell and Tissue Core at the University of Pittsburgh (Control and COPD lung) was homogenized in Trizol, total RNA was isolated according to the manufacturer’s instructions (Thermo Fisher, Grand Island, NY) and RNA was reverse-transcribed (Applied Biosystems, Grand Island, NY). Real time PCR was performed using total cDNA and primer pairs flanking introns specifically targeted the genes of interest (*slc25a4*, *slc25a5*, and *ACTB*) using Sybr green master mix for real time PCR (Applied Biosystems, Grand Island, NY). Real time PCR primer pairs included *slc25a4*, forward 5’-TGG ATG ATT GCC CAG AGT GT and reverse 5’-GGC TCC TTC GTC TTT TGC AA-3’; *slc25a5*, forward 5’-GGC TTT AAC GTG TCT GTG CA-3’ and reverse 5’-ATA GGA AGT CAA CCC GGC AA-3’; *slc25a6*, forward 5’-ACG CCC TCC ATT CAC TCT C -3’ and reverse 5’-GCT TGA CCC GCT CGA TCG -3’; *slc25a31*, forward 5’-TTC CGC TTC CCT TCA TCG TA-3’ and reverse 5’-GCC ACC GCT GTC TTG GAC-3’; *ACTB*, forward 5’-ATC CGC CGC CCG TCC-3’ and reverse 5’-CGA TGG AGG GGA AGA CGG-3’). Each sample was measured in quadruplicate. A single technical outlier was removed from samples with technical SD greater than 1, individual samples were excluded if removal of an outlier failed to reduce technical SD to less than 1 in either the gene of interest or the housekeeping gene. Relative fold change was calculated by normalizing to beta-actin and comparing between lung tissue from individuals with a history of COPD and those without a history of COPD using the ∆∆CT method. Error is calculated as positive or negative error = 2^(Fold change ± √(SEM of *ACTB*^2 + SEM of gene^2)).

For mouse lung tissues, total RNA was extracted using Trizol with analysis by real time PCR according to manufacturer’s instructions (One-step VERSO SYBR green PCR kit, Thermo Fisher Scientific). Total RNA was extracted from mouse lung tissue using Trizol with analysis of 25 ng of RNA reverse transcribed and complementary DNA was amplified by real time-PCR using a One-step VERSO SYBR green PCR kit (Thermo Fisher Scientific) on a BioRad CFX96 Real Time PCR machine. The PCR protocol included cDNA synthesis 50°C for 15 min followed by repeat cycles of inactivation at 95°C for 15 min, denaturing of DNA at 95°C for 15 seconds, annealing at 60°C for 30 seconds, extension at 72°C for 30 seconds (repeat for a total of 40 cycles). PCR primer efficiency was determined, amplicon melting curves (60 to 95°C and single product of correct size by gel analysis were determined to verify production of single amplicons. No template and no enzyme controls were compared. Target primers were used at 200 nM and include: Sigma KiQ primer sets for mouse *slc25a4*/ANT1 (#M_Slc25a4_1) and mouse *slc25a5*/ANT2 (#M_Slc25a5_1); Other primers include mouse GAPDH, human *slc25a4*/ANT1 (forward 5’-TGG ATG ATT GCC CAG AGT GT; reverse 5’-GGC TCC TTC GTC TTT TGC AA). The real time PCR data were analyzed by the ∆∆CT method with normalization to GAPDH and comparing smoke exposed mouse lung tissue to air-exposed controls.

#### Human Microarray Data

Microarray data was analyzed from the Lung Genome Research Consortium (LGRC, University of Pittsburgh) and using the publicly available Geo Omnibus Data set, GDS2486 previously published [45, 46]. The LGRC data set is a genome-wide association study of human whole lung tissue mRNA from control individuals (n=137) and those with COPD (n=219). The Geo Omnibus Data set GDS2486, previously published [45, 46] and publicly available, includes Affymetrix analysis (Affymetrix Human Genome U133 Plus 2.0 Array) of human small airway epithelial cell brushings from non-smokers (n=12) versus smokers (n=10). Data were analyzed *a priori* for gene expression of *SLC25A4*/ANT1 and *SLC25A5*/ANT2 with normalization to GPI (glucose-6-phosphate isomerase). Gene expression data do not represent multivariate comparisons and statistical analysis was completed using a Student’s t-test with p<0.05 considered to be statistically significant.

#### Human Protein Expression

Immunoblot analysis was completed on cell culture protein lysates, yeast *S. cerevisiae* protein lysates, or mouse lung tissue homogenates. Mammalian cell culture lysates were attained using RIPA buffer with protease inhibitor cocktails I, II and III (Sigma), RNase, and 150 nM aprotinin. Yeast protein lysates were obtained from yeast expressing human ANT1-4 (∆*aac[EV],* ∆*aac[ANT1],* ∆*aac[ANT2],* ∆*aac[ANT3],* ∆*aac[ANT4],* OD_600_=3 per group) via alkaline lysis with NaOH/β-mercaptoethanol and trichloroacetic acid. Protein concentration was determined by Bradford Assay (Pierce). Proteins were resolved by 10-15% SDS-polyacrylamide gel electrophoresis and transferred to a nitrocellulose membrane. Proteins of interest were immunoblotted using primary antibodies (incubated overnight at 4°C) followed by detection with Li-Cor secondary fluorophore conjugated antibodies using a Li-Cor Odyssey CLx. GAPDH and Ponceau S membrane staining were used as protein loading controls. Antibodies that recognize the following proteins were used (dilutions, source, and species are listed in parentheses): human ANT1 (1:500, ab1F3H11 made by lab of S. Claypool (*17*), mouse), ANT2 (1:500, ab5H7 from S. Claypool (*47*), mouse), TOM20 (1:1000, Abcam #ab186734, rabbit monoclonal), GAPDH (1:1000, Life Technologies A6455, rabbit). Antibody 1F3H11 for ANT1 (paralog specific) only worked for western analysis.

#### Human/Mouse Immunohistochemistry and Immunocytochemistry

Mouse lungs were inflation fixed with 10% buffered formalin for 24 hr and embedded in paraffin for sectioning. Fresh frozen and formalin fixed paraffin embedded human lungs were also stained from normal case controls and COPD patients. Sections were cut at 5-7 μm and were adhered to slides for 60 min at 60°C followed by deparaffinizing with xylene and rehydration in an ethanol series (for paraffin tissues). The sections were treated with sodium citrate buffer at 95°C for antigen retrieval. Human and mouse lung sections were stained for colocalization with primary and secondary antibodies: ANT1 (Abcam #ab102032 rabbit polyclonal, 1:1000), pan ANT (anti-ANT1/2/3 Abcam, #ab110322, mouse monoclonal 1:100), ANT2 (5H7, from S. Claypool, 1:100), ANT2 (Abcam, #ab118076, mouse monoclonal, 1:100), ANT2 (Abcam, #ab222843, 1:1000), ANT2/3 (Abcam, #ab230545, 1:100), acetylated α-tubulin (Cell Signaling, #5335T Rabbit at 1:2000 and Invitrogen, #32-2700 Mouse at 1:2000), TOM20 (Abcam, #ab56783 and Cell signaling, #42406S), NPHP4 (Atlas Antibodies, #HPA065526), GFP (Aves Labs Inc, 1:1000), goat anti-mouse, anti-rabbit or anti-chicken Alexa 488, 555 and 647 (Molecular Probes). Antibodies used for each experiment are designated in the figures. Images were captured on a Zeiss 780 confocal microscope with an iPlan Apochromat 63x/1.4 NA oil objective, on a Nikon A1R with 60x objective or Nikon SIM microscope using a 100x objective with SIM reconstruction after image capture. Control sections were stained with non-immune rabbit IgG (#2729P, Cell signaling). Quantification of ANT staining was determined by making regions of interest [2] of the ciliary region with a second ROI with the associated mitochondrial area below that layer (see **supplementary Fig. S5B**, **S7B**). Approximately 5-14 measurements were made per ciliated area of epithelium per image. Mean pixel intensities were normalized to background by subtracting the mean pixel intensity for a large background ROI. The sum was calculated separately for the ROI areas and ROI normalized mean intensities for the ciliary and mitochondrial areas. The integrated intensity was calculated by dividing the sum of the ROI normalized mean intensities by the sum of the ROI areas. Images with multiple colors are displayed with each channel in gray scale for clarity and a composite pseudocolor image.

For immunocytochemistry on cells in culture on glass or on inserts at air liquid interface, cells were fixed in fresh 4% paraformaldehyde for 10 min at 4°C. For select experiments to label mitochondria, cells were incubated with Mitotracker Deep Red (Invitrogen, #M22426) for 20 min at 37°C according to manufacture protocol followed by PFA fixation. Cells were washed three times in PBS, permeabilized with ice cold 0.3% Triton X-100 with 1% BSA in PBS for 10 min followed by three washes in PBS. To prevent non-specific staining, cells were blocked with 2% BSA in PBS for 45 min at room temperature. Cells were washed five times with 0.5% BSA in PBS. Cells were incubated with primary antibody overnight at 4°C followed by five washes with 0.5% BSA in PBS. Cells were incubated with secondary antibody dilutions (made in 0.5% BSA in PBS) for 60 min at room temperature in the dark. Cells are washed 5 times with 0.5% BSA in PBS followed by five PBS washes. Nuclei were stained with Hoechst at 10 μg/mL for 10 min followed by PBS washes. For insert staining, insert membranes are cut out and placed apical surface upright onto glass slides. Sections were mounted in Prolong Diamond Antifade Mounting Agent (Molecular Probes, ThermoFisher), cured for at least 24 hr at room temperature, and sealed with clear nail polish. Samples were stored at 4°C protected from light for long-term storage. To assess the membrane localization of ANTs using differential permeabilization, ALI cultured cells were blocked as above, incubated with the noted primary antibody against ANT with subsequent washes, followed by 1 hr incubation with an Alexa 555 secondary antibody all without permeabilization. Then, the tissues were permeabilized and incubated again with the same primary antibody against ANT for 2 hr at room temp, washed, and incubated for 1 hr with an Alexa 488 secondary antibody. The remaining processing was similar for subsequent confocal imaging.

#### Molecular Phylogenetic Tree Analysis

The evolutionary history was inferred by using the Maximum Likelihood method based on the JTT matrix-based model [47]. The bootstrap consensus tree inferred from 500 replicates [48] is taken to represent the evolutionary history of the taxa analyzed. Branches corresponding to partitions reproduced in less than 50% bootstrap replicates are collapsed. The percentage of replicate trees in which the associated taxa clustered together in the bootstrap test (500 replicates) are shown in red next to the branches [48]. Initial tree(s) for the heuristic search were obtained automatically by applying Neighbor-Join and BioNJ algorithms to a matrix of pairwise distances estimated using a JTT model, and then selecting the topology with superior log likelihood value. The tree is drawn to scale, with branch lengths measured in the number of substitutions per site. The analysis involved 22 amino acid sequences. All positions containing gaps and missing data were eliminated. There was a total of 289 positions in the final dataset. Evolutionary analyses were conducted in MEGA7 [49].

#### Quantification and Statistical Analysis

Mean densitometry and all other quantitative data (mean ± SEM) are normalized to appropriate control groups and assessed for normalcy. If the samples are normally distributed, then the data sets are analyzed for significance using ANOVA followed by a Fisher’s LSD post-test. If the data are not normally distributed, nonparametric analyses, including Kruskal-Wallis and/or Mann-Whitney are used.

## Supporting information

Kliment et al. Supplemental Data

## Acknowledgements

We thank the University of Pittsburgh Lung Tissue and Airway Cell Core funded by the CF Foundation Research Development Grant (J. Pilewski and M. Myerburg), Randal Reed (Johns Hopkins University) for mouse nasal epithelial tissue sections, Carolyn Machamer (Johns Hopkins University) for helpful discussions and Corinne Sandone (Johns Hopkins University) for the artwork in Figure 1 and 7. We thank members of the Robinson lab (Johns Hopkins) and Steve Shapiro (University of Pittsburgh) for helpful comments on the manuscript. We would like to thank Dr. Douglas Wallace, Children’s Hospital of Pennsylvania for the ANT1 knockout mice. We thank the University of Pittsburgh Center for Biological Imaging (S. Watkins and C. St. Croix) and the Johns Hopkins Institute for Basic Biomedical Sciences Microscope Facility for imaging assistance (Confocal Instrument grant NIH S10OD016374).

## Author Contributions

CRK and DNR designed experiments. CRK performed the cDNA library selection and *Dictyostelium* growth studies, AncA plasmid cloning, adenovirus characterization, mammalian cell viability studies, immunohistochemistry and immunocytochemistry, Vitrocell Smoke Exposure System experiments, lentiviral primary cell studies including ASL, cell imaging experiments and quantitative PCR. CRK and JMN performed airway surface liquid imaging and ciliary beat frequency studies. JMN performed MitoSOX oxidative stress studies. CRK, MJK and JMN analyzed data. PAI developed the MatLab analysis script for ASL and CBF assessments. VKS provided experimental input and LTRC Lung samples. YWL and SMC provided ANT antibodies, yeast expressing human ANT, metabolic Seahorse study input, and experimental design guidance. ADG performed and completed the mouse smoke exposure studies and lung sample harvest. FCS, JER and YZ provided and analyzed the human lung tissue for PCR gene expression and the LGRC data. CRK and DNR wrote the manuscript, and all authors participated in editing.

## Data Availability

All data associated with this study are available in the main text or the supplemental materials. All data and analytical codes are available from the corresponding authors upon reasonable request.

## SUPPLEMENTAL MATERIALS

Supplemental materials include nine Supplementary Figures S1-S9 and eight Supplementary Videos 1-8.

### Supplementary Figures

Supplementary Fig. S1. ANT is a genetic protector of *Dictyostelium* and bronchial epithelial cells from cell death.

Supplementary Fig. S2. Alterations in mitochondrial reactive oxygen species and metabolism due to ANT.

Supplementary Fig. S3. Airway surface liquid and ciliary beat frequency analysis.

Supplementary Fig. S4. ANT localization in the human airway epithelium.

Supplementary Fig. S5. ANT antibody panel in the ciliated airway epithelium of normal human lung tissue.

Supplementary Fig. S6. ANT antibody specificity by Western blot analysis.

Supplementary Fig. S7. ANT antibody panel in ciliated human bronchial epithelial cells grown at air liquid interface.

Supplementary Fig. S8. Method for airway surface liquid (ASL) thickness.

Supplementary Fig. S9. Method for computing ciliary beating frequency (CBF).

### Supplemental Video Legends

**Supplementary Video 1.**

Kliment et al. Suppl Video 1_CTRL pre-CS

Ciliary beat frequency (CBF) in NHBE cells (control adenovirus) before exposure to CS.

**Supplementary Video 2.**

Kliment et al. Suppl Video 2_CTRL post-CS

CBF in NHBE cells (control adenovirus) 30 min after exposure to CS, showing a slowing of CBF.

**Supplementary Video 3.**

Kliment et al. Suppl Video 3_ANT1 pre-CS

CBF in NHBE cells (ANT1-GFP adenovirus) before exposure to CS.

**Supplementary Video 4.**

Kliment et al. Suppl Video 4_ ANT1 post-CS

CBF in NHBE cells (ANT1-GFP adenovirus) 30 min after exposure to CS, showing a slowing of CBF.

**Supplementary Video 5.**

Kliment et al. Suppl Video 5_ ANT2 pre-CS

CBF in NHBE cells (ANT2-GFP adenovirus) before exposure to CS.

**Supplementary Video 6.**

Kliment et al. Suppl Video 6_ ANT2 post-CS

CBF in NHBE cells (ANT2-GFP adenovirus) 30 min after exposure to CS, showing preservation of CBF.

**Supplementary Video 7.**

Kliment et al. Suppl Video 7_Air treated mouse lung

SIM imaging of air-treated mouse lung stained for ANT1 (red, Rabbit anti-human ANT1 antibody, ab102032), tubulin (light blue) and nuclei (dark blue). ANT1 localizes to the plasma membrane surface in a linear pattern on the apical cell surface and to cilia in a striated pattern. ANT also localizes to mitochondria in the cell body.

**Supplementary Video 8.**

Kliment et al. Suppl Video 8_Smoke treated mouse lung

SIM imaging of cigarette smoke-treated mouse lung stained for ANT1 (red, rabbit anti-human ANT1, ab102032), tubulin (light blue) and nuclei (dark blue). ANT1 localizes to the plasma membrane surface in a linear pattern on the apical cell surface and to cilia with loss of patterning. ANT also localizes to mitochondria in the cell body.

## Key Resources

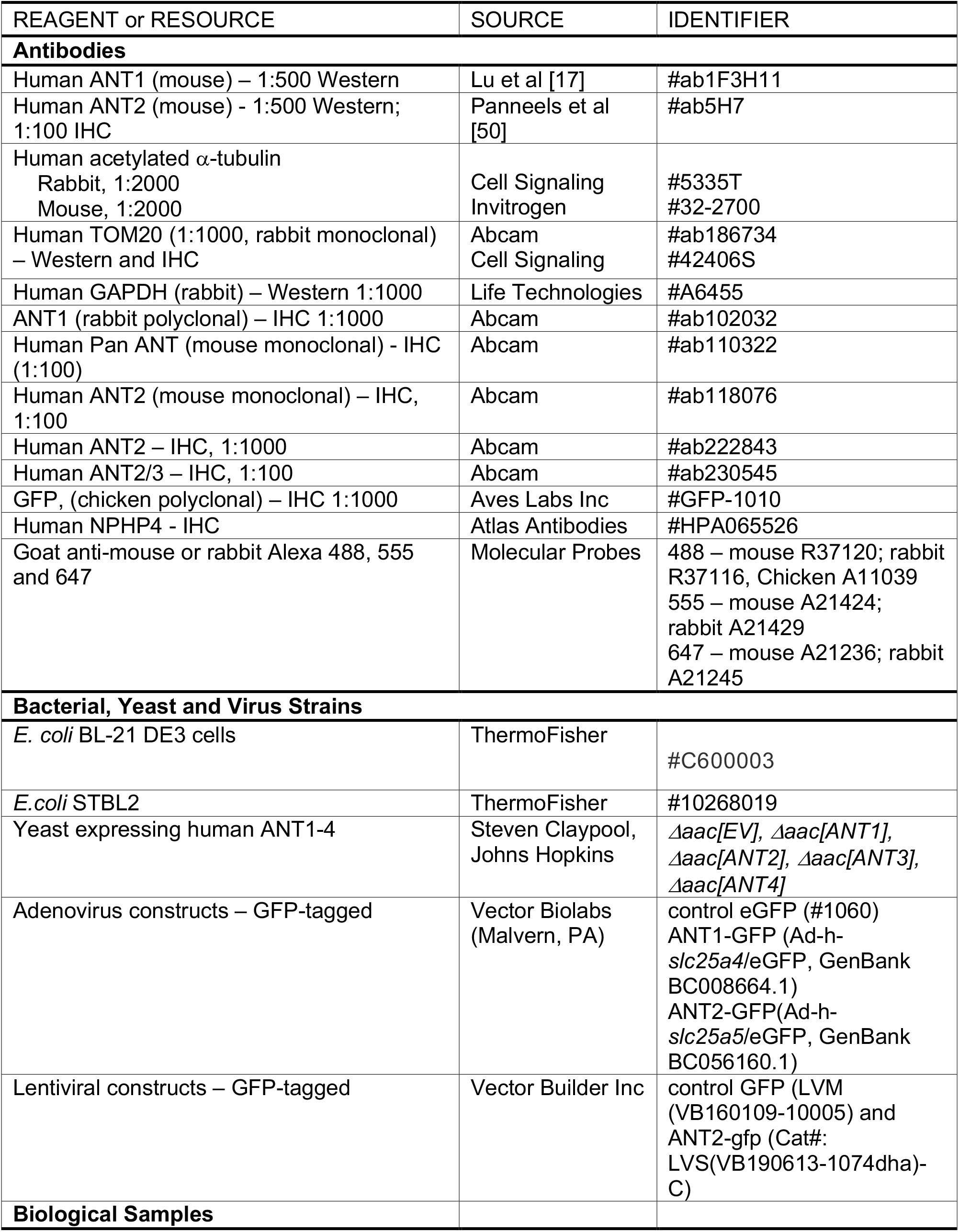
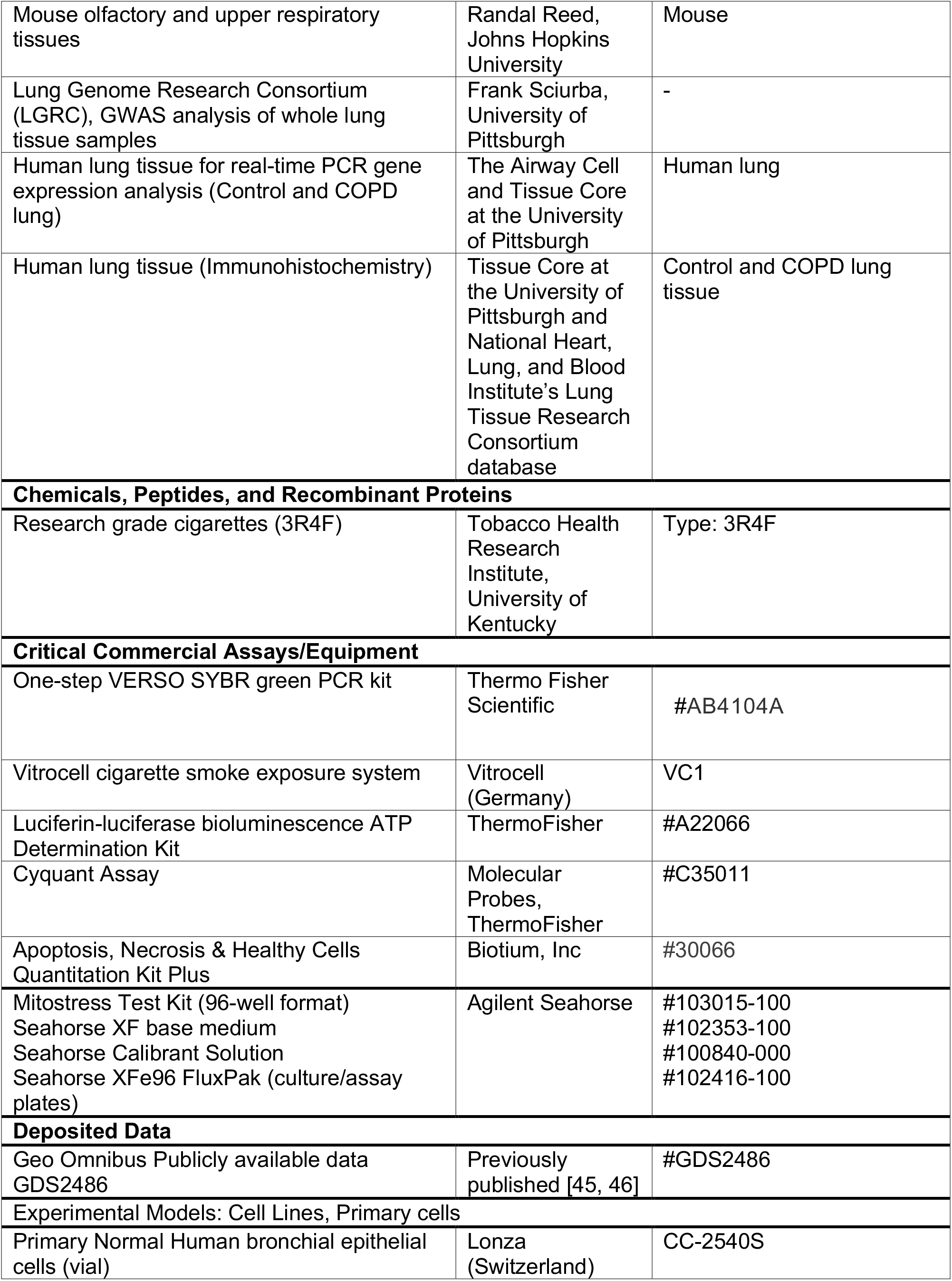
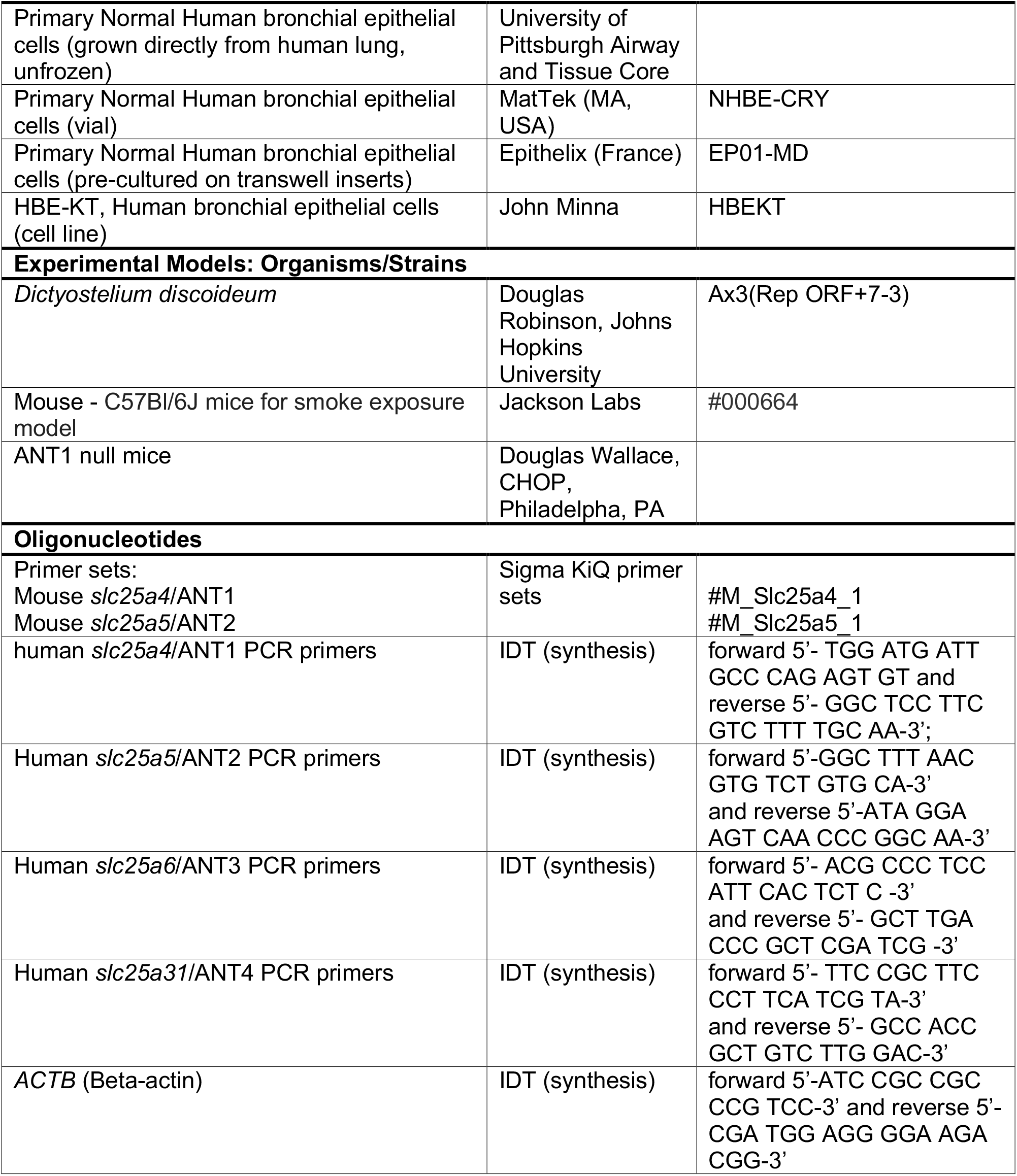
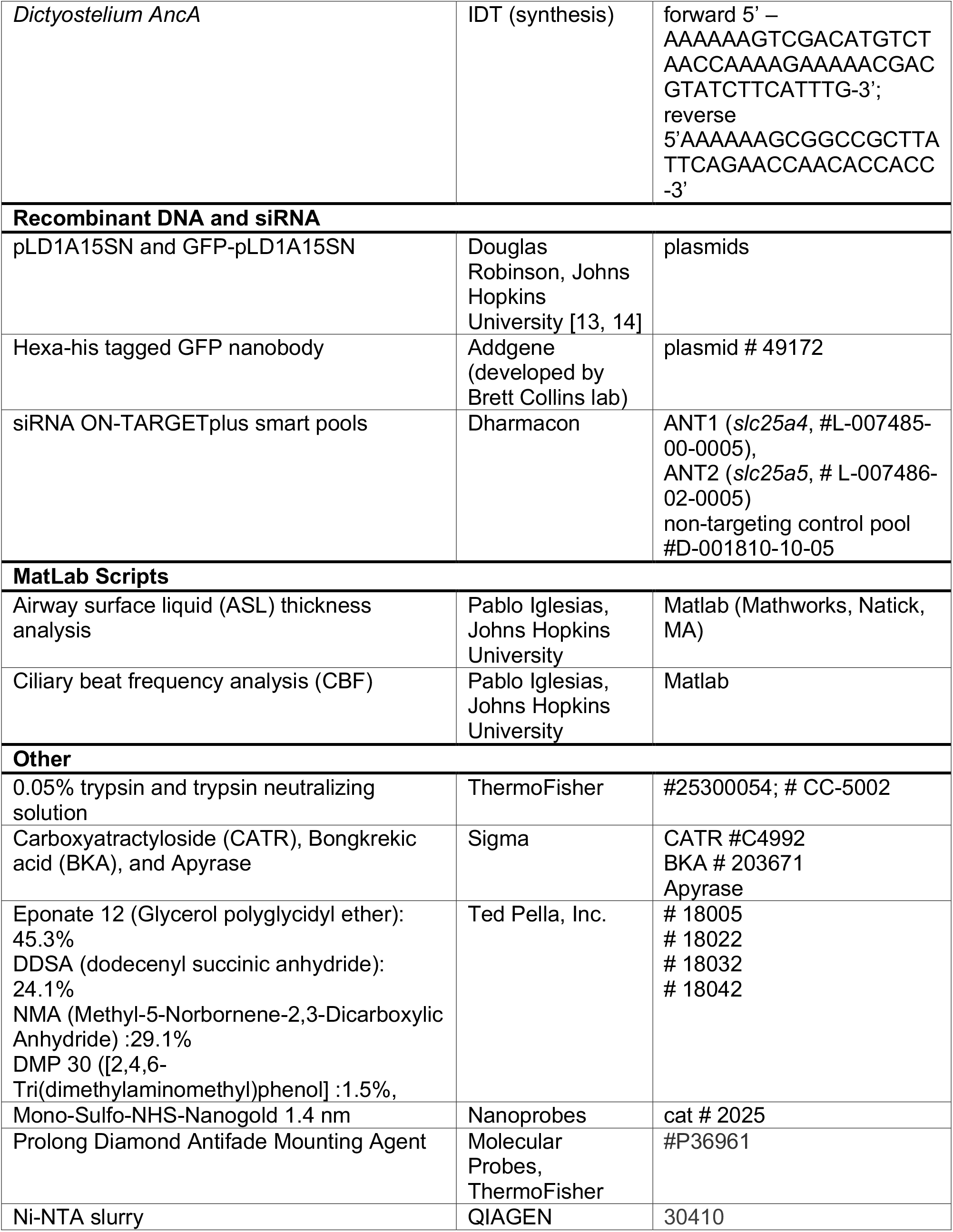
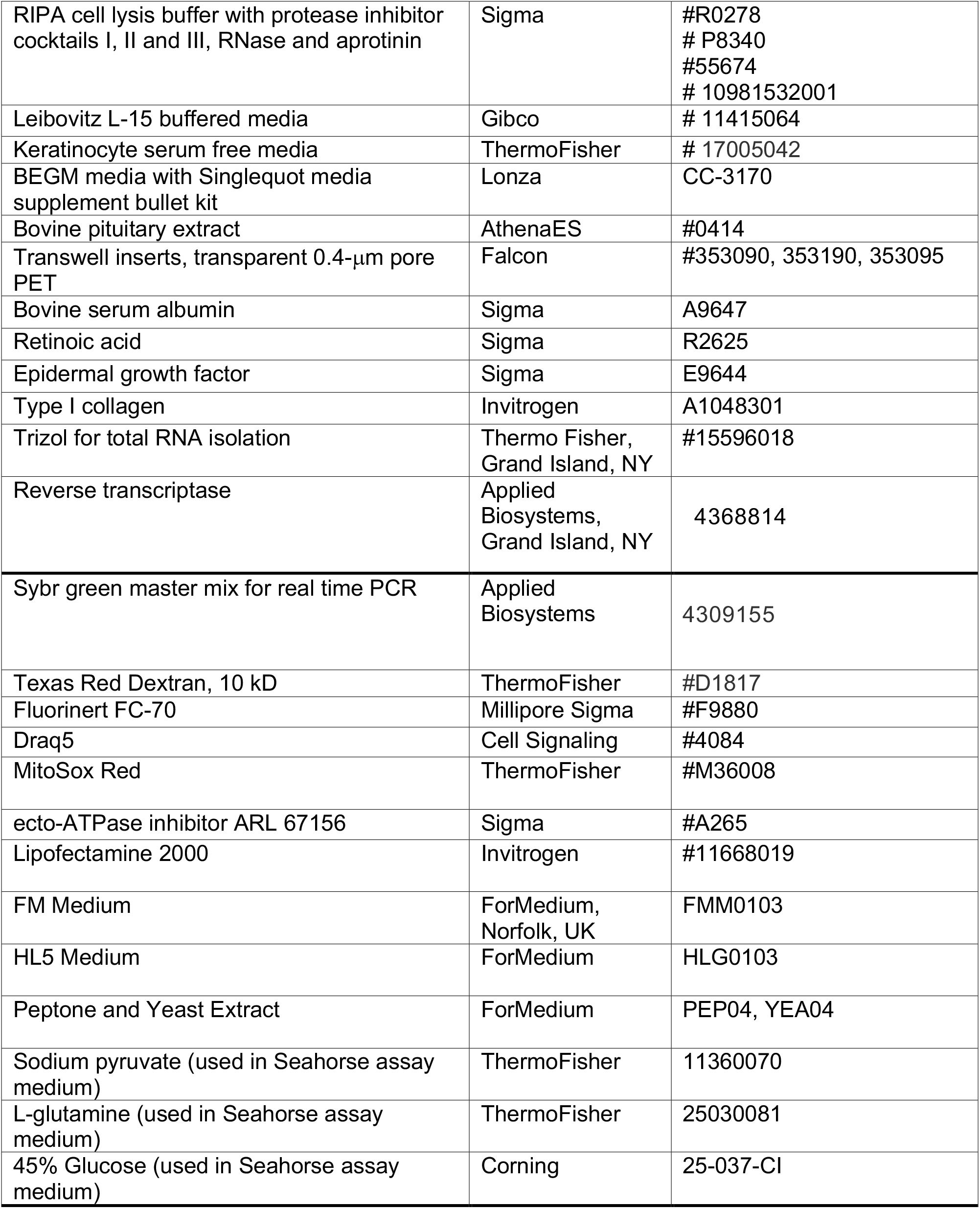

