## Supplementary material for "Adenine Nucleotide Translocase regulates the airway epithelium, mitochondrial metabolism and ciliary function": Kliment et al. Supplemental Data

### Supplemental Materials

Materials include Supplemental Figures S1 to S9 and Legends to Supplemental Videos 1-8.

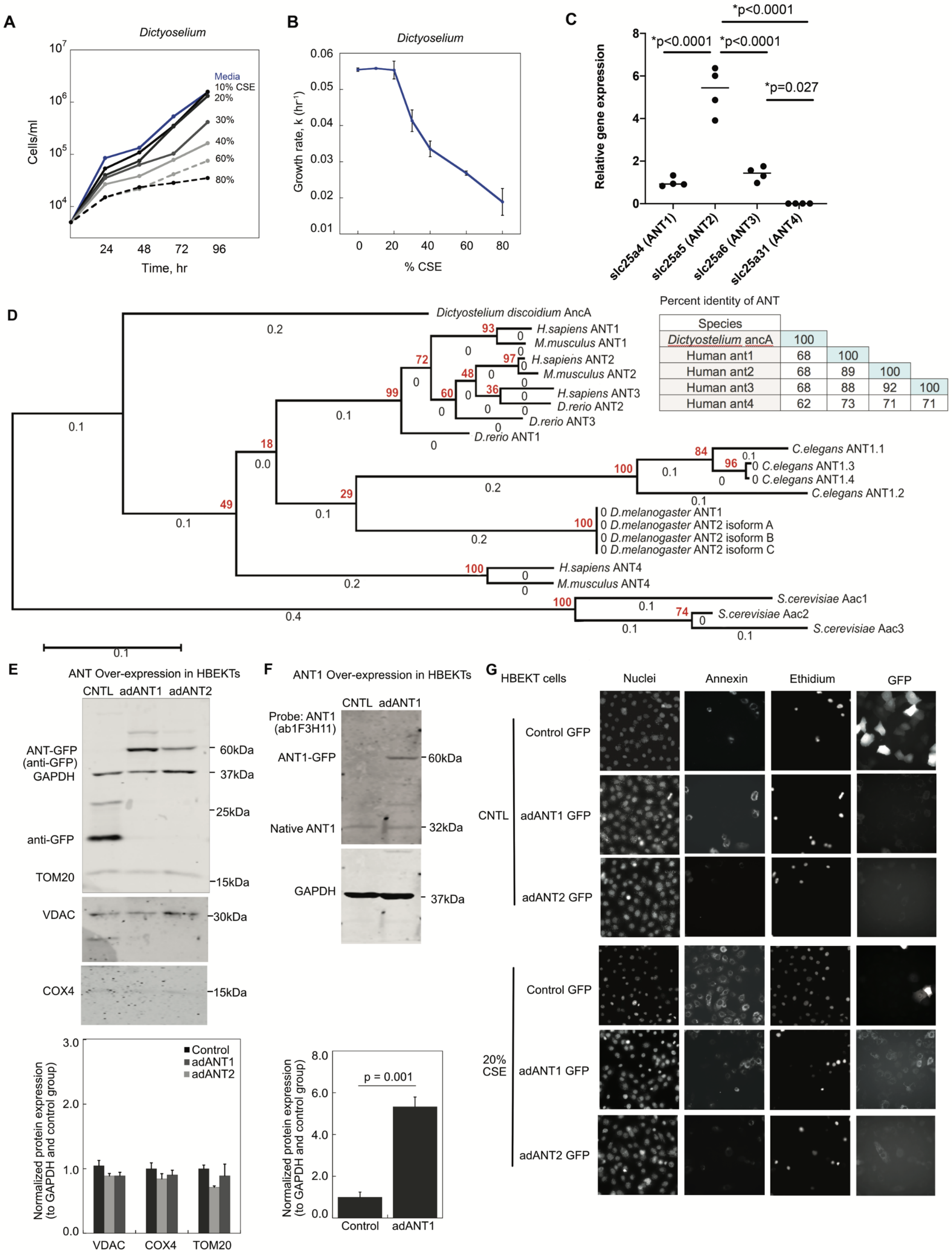


| Species | | | | |
|---|---|---|---|---|
| Dictyostelium ancA | 100 | | | |
| Human ant1 | 68 | 100 | | |
| Human ant2 | 68 | 89 | 100 | |
| Human ant3 | 68 | 88 | 92 | 100 |
| Human ant4 | 62 | 73 | 71 | 71 |

**Supplementary Fig. S1. ANT is a genetic protector of *Dictyostelium* and bronchial epithelial cells from cell death. A)** *Dictyostelium* growth curves with 10-80% CSE. **B)**

*Dictyostelium* growth curve to identify EC40 for 10-80% CSE. **C)** Relative gene expression of human *slc25a4* (ANT1), *slc25a5* (ANT2), *slc25a6* (ANT3) and *slc25a31* (ANT4) in primary differentiated normal human bronchial epithelial cells grown at air liquid interface. n = 4 inserts. Statistics by Mann Whitney; p-values are noted. **D)** Molecular phylogenetic tree analysis of adenine nucleotide translocase (AncA in *Dictyostelium* and ANT in humans) with bootstrap values. Percent identity for human ANT and *Dictyostelium* AncA, generated by Clustal2.1. **E)** Western analysis of adenoviral overexpression (control GFP, ANT1-GFP and ANT2-GFP) in HBEKTs evaluating for GFP, mitochondrial TOM20, VDAC, COX4 and GAPDH. **F)** ANT1 western analysis for ANT1 overexpression. **G)** Representative images of HBEKTs depicting total nuclei (Draq5), apoptosis (Annexin V), necrosis (ethidium homodimer) and GFP (adenovirus infection).

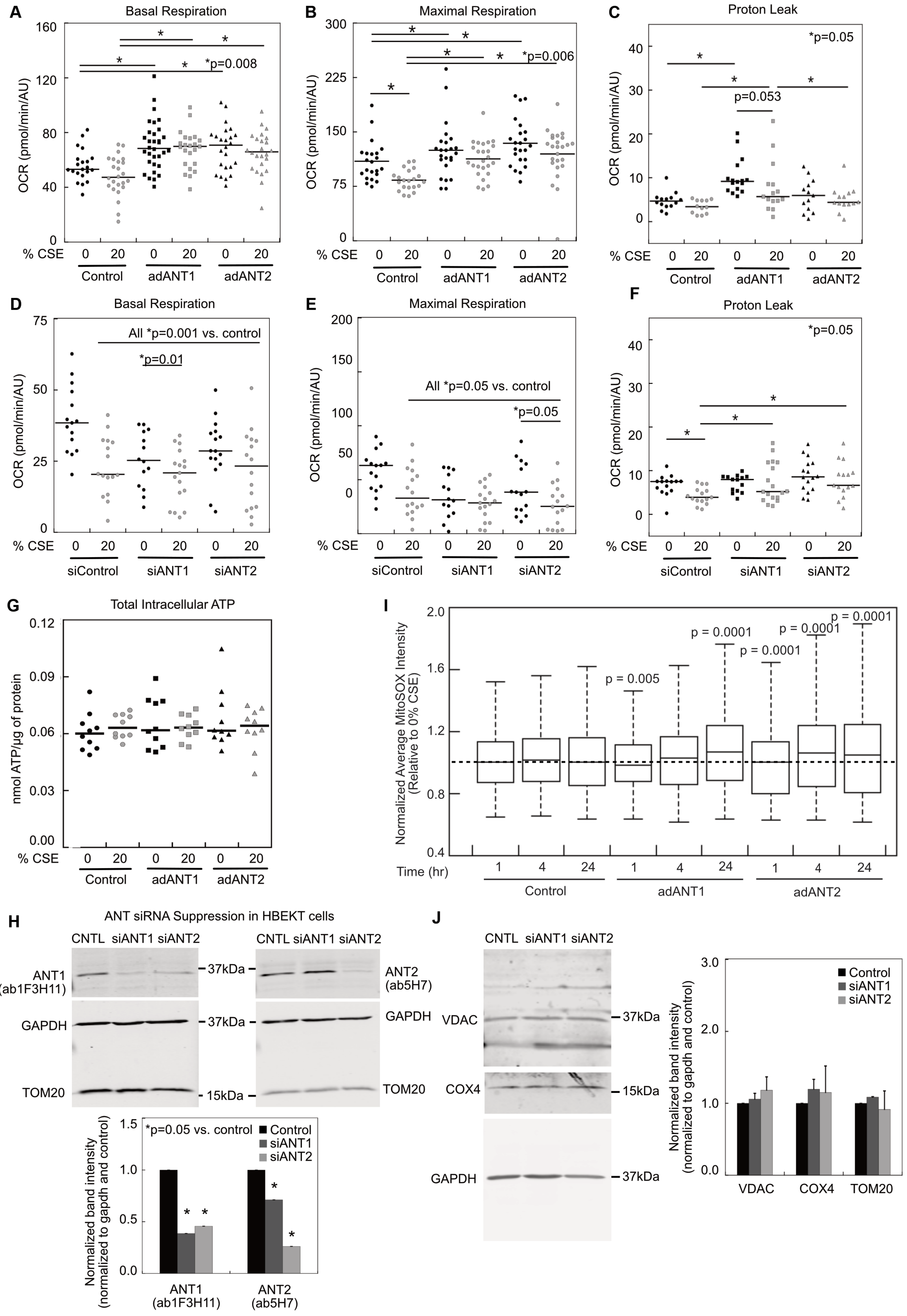
A
Basal Respiration
OCR (pmol/min/AU)
*p=0.008
% CSE 0 20 0 20 0 20
Control adANT1 adANT2
B
Maximal Respiration
OCR (pmol/min/AU)
*p=0.006
% CSE 0 20 0 20 0 20
Control adANT1 adANT2
C
Proton Leak
*p=0.05
p=0.053
OCR (pmol/min/AU)
% CSE 0 20 0 20 0 20
Control adANT1 adANT2
D
Basal Respiration
All *p=0.001 vs. control
*p=0.01
OCR (pmol/min/AU)
% CSE 0 20 0 20 0 20
siControl siANT1 siANT2
E
Maximal Respiration
All *p=0.05 vs. control
*p=0.05
OCR (pmol/min/AU)
% CSE 0 20 0 20 0 20
siControl siANT1 siANT2
F
Proton Leak
*p=0.05
OCR (pmol/min/AU)
% CSE 0 20 0 20 0 20
siControl siANT1 siANT2
G
Total Intracellular ATP
nmol ATP/µg of protein
% CSE 0 20 0 20 0 20
Control adANT1 adANT2
I
Normalized Average MitoSOX Intensity (Relative to 0% CSE)
p = 0.005
p = 0.0001
p = 0.0001
p = 0.0001
p = 0.0001
Time (hr) 1 4 24 1 4 24 1 4 24
Control adANT1 adANT2
H
ANT siRNA Suppression in HBEKT cells
CNTL siANT1 siANT2
CNTL siANT1 siANT2
ANT1 (ab1F3H11)
ANT2 (ab5H7)
GAPDH
TOM20
37kDa
15kDa
*p=0.05 vs. control
Control
siANT1
siANT2
Normalized band intensity (normalized to gapdh and control)
ANT1 (ab1F3H11)
ANT2 (ab5H7)
J
CNTL siANT1 siANT2
VDAC
COX4
GAPDH
37kDa
15kDa
Control
siANT1
siANT2
Normalized band intensity (normalized to gapdh and control)
VDAC COX4 TOM20

**Supplementary Fig. S2. Alterations in mitochondrial reactive oxygen species and metabolism due to ANT.** Cellular metabolism by the Seahorse Mitostress assay in HBEKT cells, with ANT1 or ANT2 adenoviral overexpression (**A**, **B**, **C**) or siRNA suppression (**D**, **E**, **F**). **A)** Basal OCR with ANT overexpression, **B)** Maximal OCR with ANT overexpression, **C)** Proton leak with ANT overexpression, **D)** Basal OCR ANT suppression, **E)** Maximal OCR with ANT suppression, **F)** Proton leak with ANT suppression, median bars are provided; n = 15-26 wells from 3 separate experiments. **G)** Measurements of total intracellular ATP in HBEKTs with ANT1 or ANT2 overexpression ±20% CSE, yielding a total intracellular [ATP] of 8 mM for HBEKT cells. Median bars shown; n = 10 wells per group from 5 separate experiments. **H)** Western analysis of ANT siRNA suppression probed for ANT1 (ab1F3H11), ANT2 (ab5H7), GAPDH and TOM20. Statistical analysis by ANOVA with Fisher's LSD posttest; *p<0.05. **I)** Boxplot showing mitochondrial superoxide production (MitoSox) in HBEKTs after CSE with ANT overexpression. Horizontal dotted line represents the median for the 1 hr control group. Boxplots show the median with the box delineating the 1$^{st}$ and 3$^{rd}$ quartiles. The whiskers represent 1.5*IQR (interquartile range). Statistics performed by Kruskal-Wallis and two-tailed Mann-Whitney U tests. P-values represent differences from Control at each respective time point. n = 2800-4500 cells per group. **J)** Western analysis of ANT siRNA suppression probed for mitochondrial proteins VDAC and COX4. A bar graph summarizes the relative amounts of each protein across samples. Values are for n = 3 per group with band intensity quantification normalized to GAPDH.

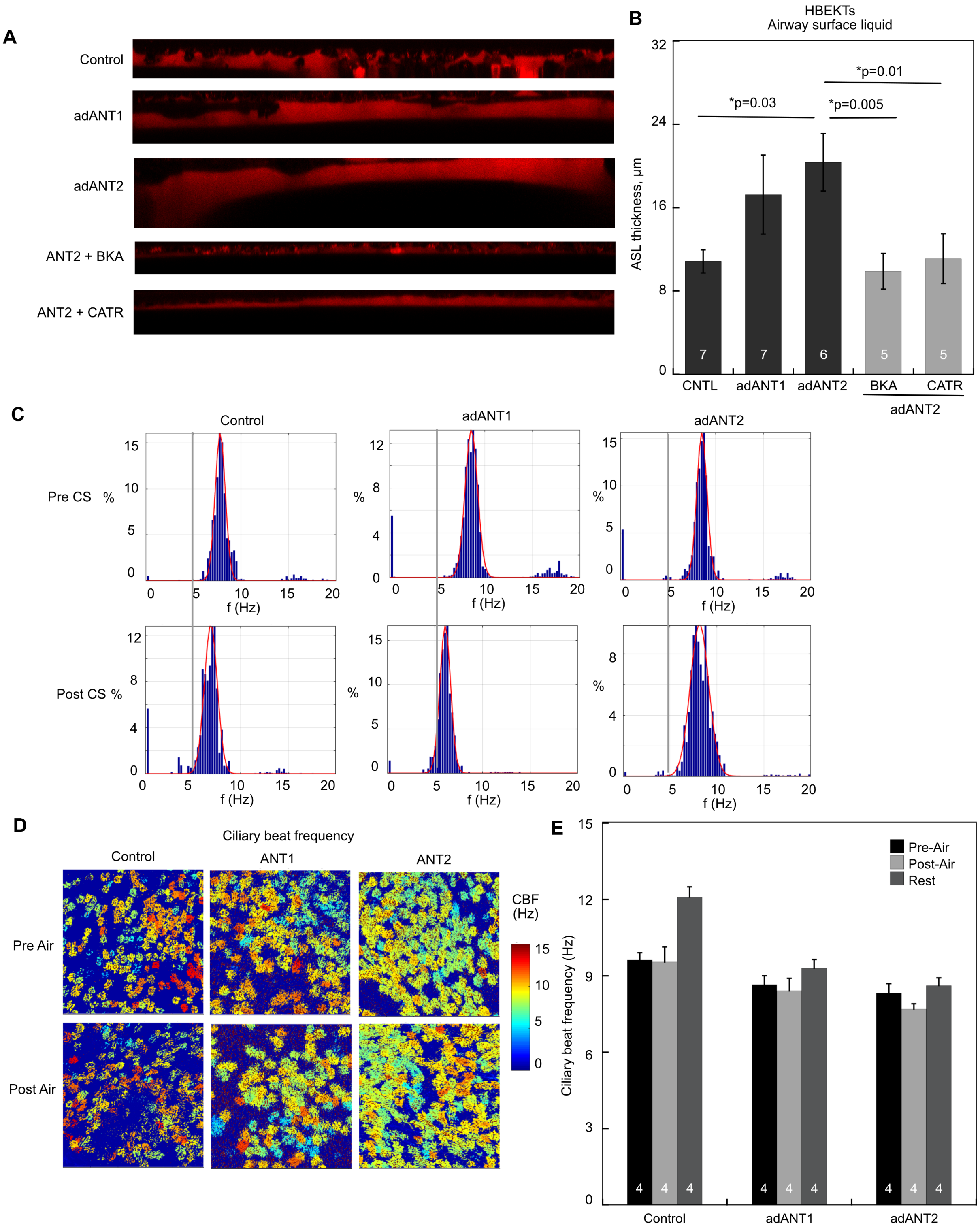


**Supplementary Fig. S3. Airway surface liquid and ciliary beat frequency analysis. A)** ASL was assessed in HBEKTs using a Texas Red dye with representative z-stack orthogonal views shown. **B)** ASL thickness for control, ANT1, or ANT2 overexpression in HBEKTs with PBS vehicle, 20 μM CATR, or 4 μM BKA treatment. **C)** CBF histograms from CS treated NHBE cells. **D)** CBF heat maps and **E)** CBF data for air-treated NHBE cells. Data show mean ± SEM and the "n" depicted on the bars equals the # of individual ALI inserts. Experiments are a compilation of insert data from 2-4 different days. Statistical analysis by ANOVA with Fisher's LSD posttest with $p<0.05$ as significant.

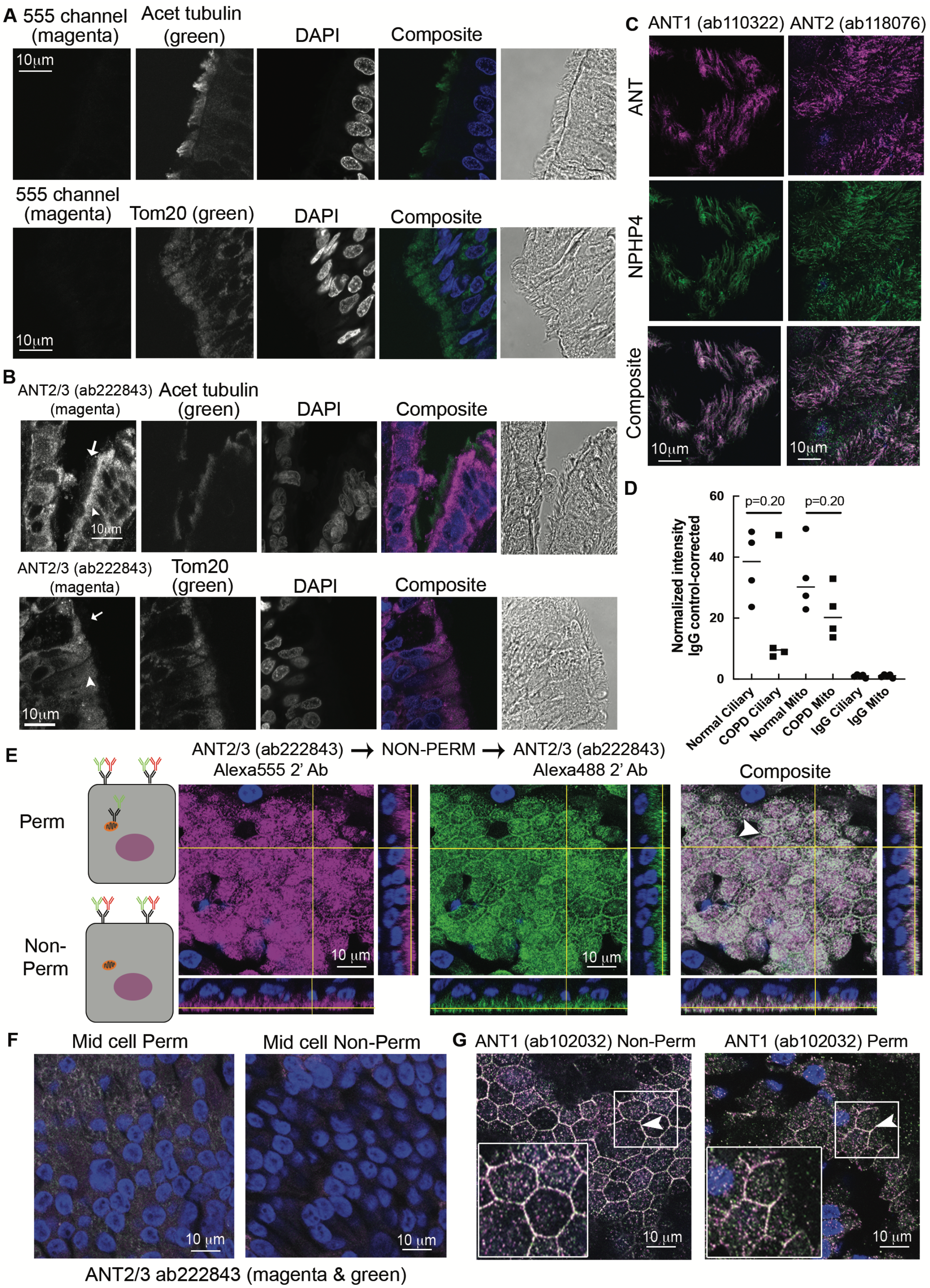


**Supplementary Fig. S4. ANT localization in the human airway epithelium. A)** Control staining of human airway tissue for acetylated $\alpha$-tubulin or TOM20 (green) without evidence of bleed through into the 555 channel (magenta). **B)** Human lung tissue with immunofluorescence for ANT2/3 (ab222843, magenta) with colocalization of ANT with acetylated $\alpha$-tubulin and TOM20 (green). The full arrow identifies the layer of cilia and arrowhead identifies the mitochondrial layer. Representative images from 2 patient tissues. **C)** Colocalization of ANT1 or ANT2 and NPHP4 (ciliary transition zone protein) in cilia of NHBEs. DAPI staining is also present, however confocal section acquired at the level of the cilia above the plane of the nucleus. **D)** Quantification of ANT staining in normal and COPD lung tissue, assessing ciliary and mitochondrial layers of staining. Corrected for background and normalized to non-immune IgG staining controls. Medians are shown. Statistical analysis using a Mann Whitney test. **E)** A population of ANT2/3 localizes to the plasma membrane in NHBE cells. NHBE cells were stained for ANT2/3 (without prior permeabilization) followed by secondary antibody Alexa 555 (magenta). Without permeabilization (non-perm), samples were subsequently stained again with anti-ANT2/3 followed by a secondary antibody Alexa 488 (green). Colocalization is evident at the cell borders and surface (white). Scale bar, 10 μm. n = 3 inserts. **F)** A representative confocal image from the middle of permeabilized and non-permeabilized cells stained with ab222843 (as in panel E) showing intracellular staining of ANT in the permeabilized ALI cells. The white inset shows additional details. **G)** ANT1 localizes to the plasma membrane in NHBE cells. In the left panel, NHBE cells were stained for ANT1 using ab102032 (secondary antibody Alexa 555, magenta) without permeabilization and subsequently stained with ab102032 (secondary antibody Alexa 488, green). In the right panel, NHBEs were stained with ab102032 (secondary Alexa 555, magenta), permeabilized (perm) and subsequently stained with ab102032 (secondary Alexa 488, green). Colocalization is depicted as white. Scale bar, 10 μm. n = 2 inserts.

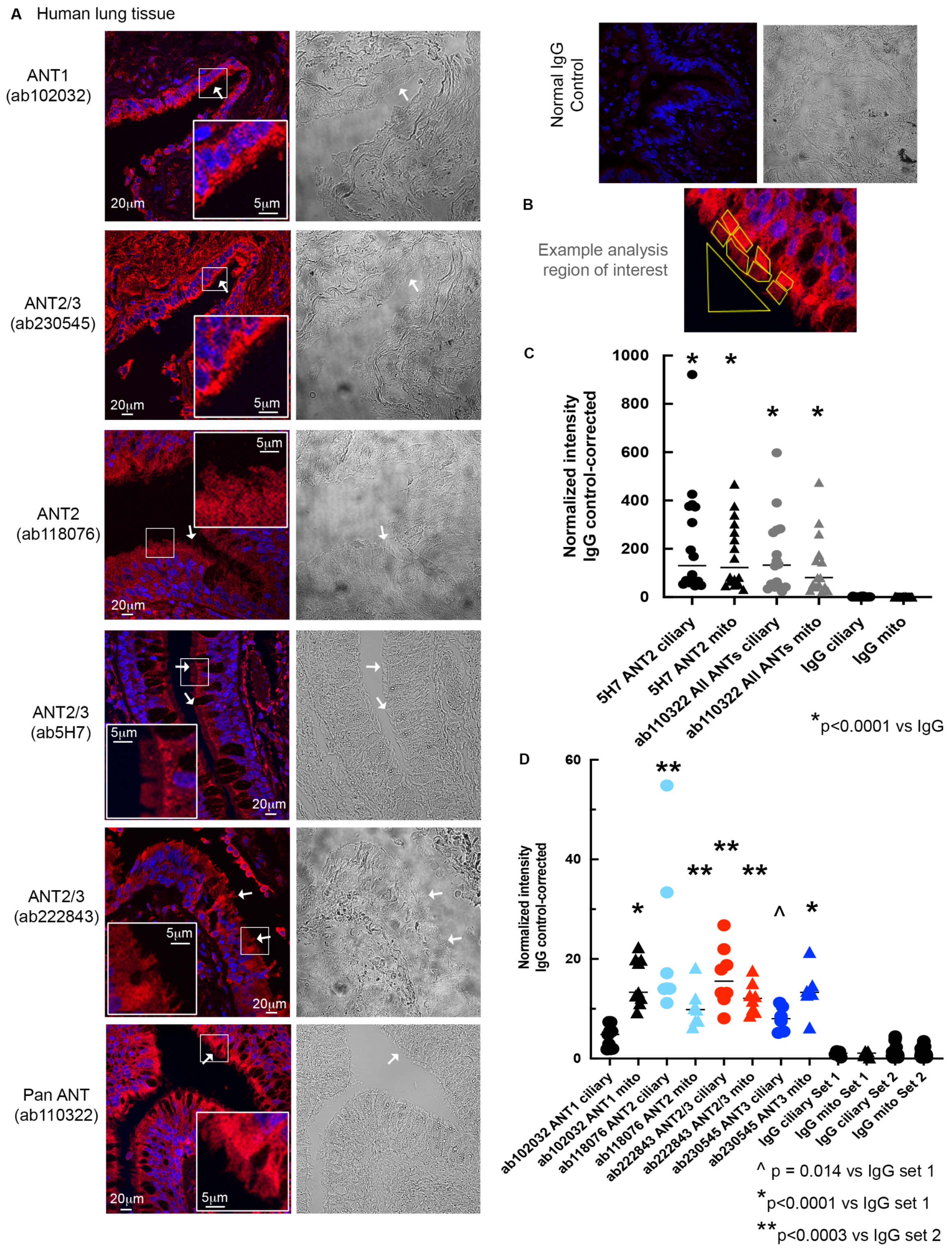


**Supplementary Fig. S5. ANT antibody panel in the ciliated airway epithelium of normal human lung tissue. A)** ANT antibodies were tested on normal human lung tissue (formalin fixed, paraffin embedded) for cellular localization. Associated DIC images are shown to allow identification of the apical layer of cilia. ANT staining at the ciliary plasma membrane is noted with the long white arrow while mitochondrial ANT is noted with the short arrow. Enlarged insets are included, scale bar 5 μm. **B)** Example of analysis completed on the apical cilia layer versus subcellular mitochondrial layer. The large triangle represents background analysis. **C)** Quantification of ciliary layer (●) versus mitochondrial layer (▲) for each antibody analyzed, ANT2 (ab5H7) and All ANTs (Pan-ANT ab110322). Each set of staining was completed with non-immune IgG staining controls which served as the normalization. n = 3 patient samples. Medians are shown. Statistics by ANOVA with Fisher's LSD posttest. p-values are noted. One patient sample was noted to have higher staining intensity as evident in the data set. **D)** Quantification of ciliary versus mitochondrial layer for each antibody analyzed, ANT1 (ab102032), ANT2 (ab118076), ANT2/3 (ab222843) and ANT3 (ab230545). Each set of staining was completed with non-immune IgG staining controls which served as the normalization. n = 3-5 patient samples. Medians are shown. Statistics by ANOVA with Fisher's LSD posttest. P-values are noted.

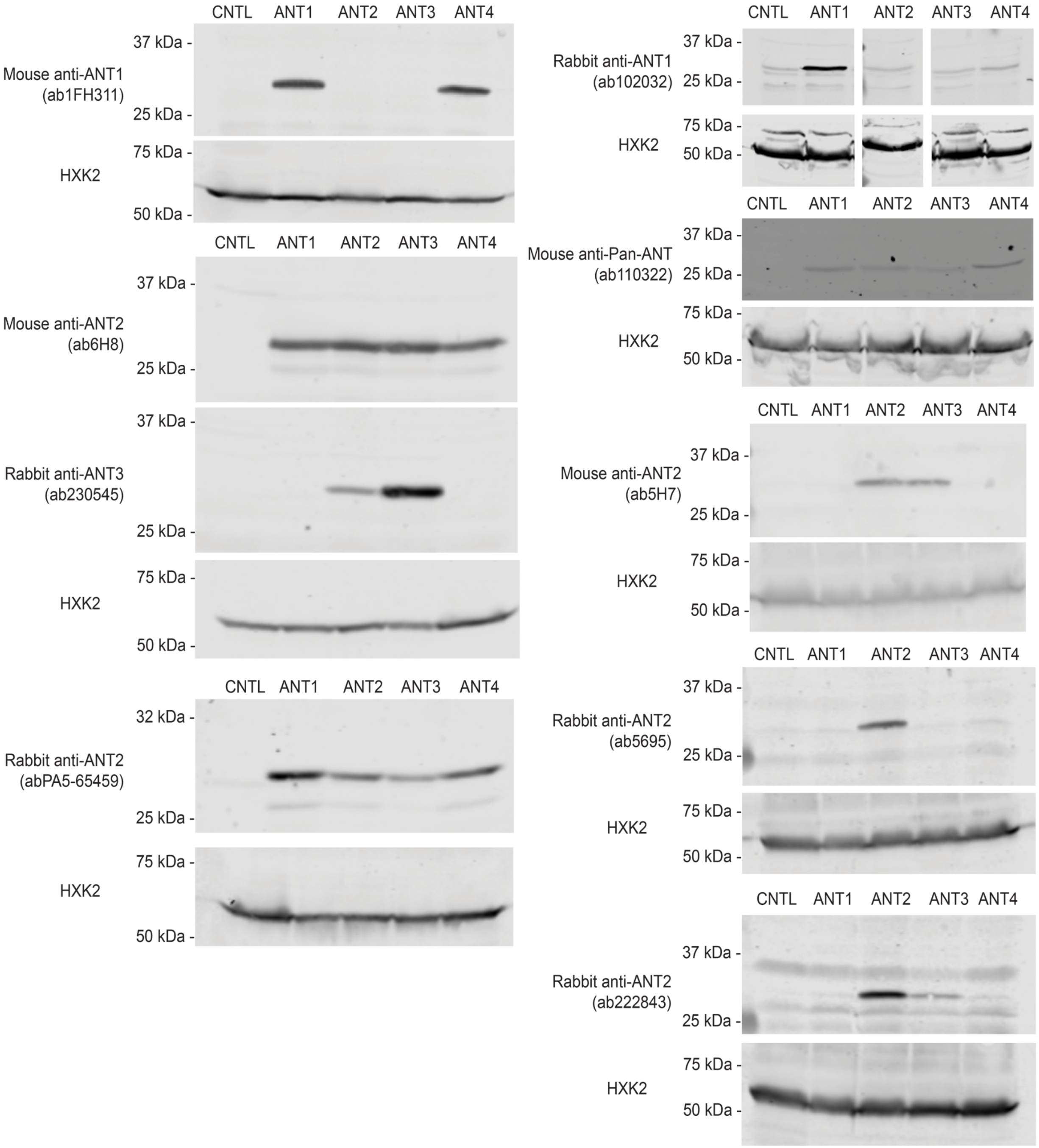


**Supplementary Fig. S6. ANT antibody specificity by Western blot analysis.** Antibody specificity was determined for rabbit and mouse anti-ANT antibodies utilizing yeast strains with expression of the individual human ANT paralogs, ANT1-4. Hexokinase-2 was used as the loading control.

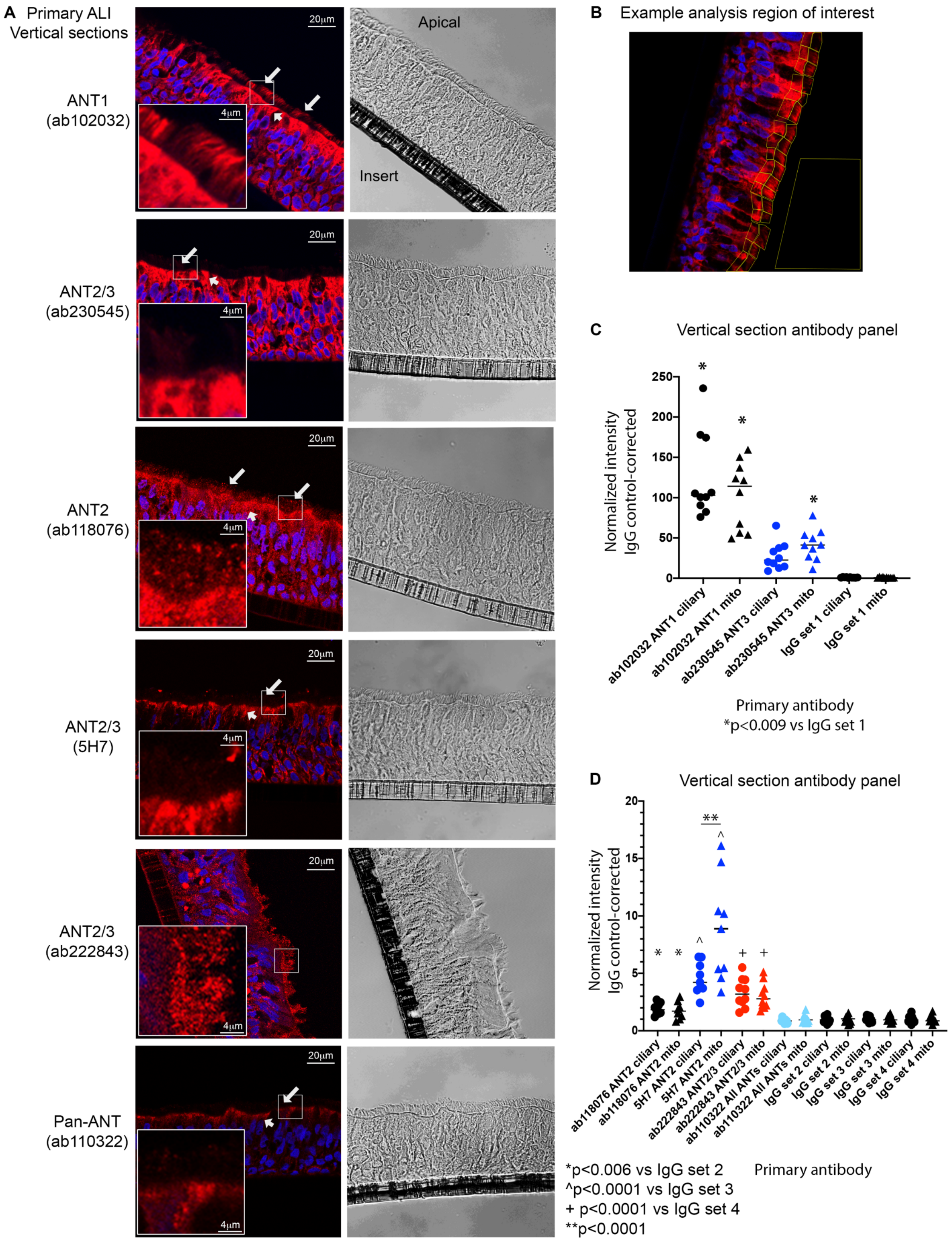


**Supplementary Fig. S7. ANT antibody panel in ciliated human bronchial epithelial cells grown at air liquid interface. A)** ANT antibodies were tested on primary normal human bronchial epithelial cells differentiated at air liquid interface on cell culture inserts. Frozen sections were cut of the cells on inserts and subsequently stained for ANT cellular localization. Associated DIC images are shown to allow identification of the apical layer of cilia. ANT staining at the ciliary plasma membrane is noted with the long white arrow while mitochondrial ANT is noted with the short arrow. Enlarged insets are included, scale bar 4 μm. **B)** Example of analysis completed on the apical cilia layer versus subcellular mitochondrial layer with an area for background analysis. Please see the immunofluorescence staining section of the methods for quantification methodology. **C, D)** Quantification of ciliary layer (●) versus mitochondrial layer (▲) for each antibody analyzed. Each set of staining was completed with non-immune IgG staining controls which served as the normalization. Represents cells from n = 3-5 patients. Median bars are shown. Statistics by ANOVA with Fisher's LSD posttest, p-values are noted in the figure.

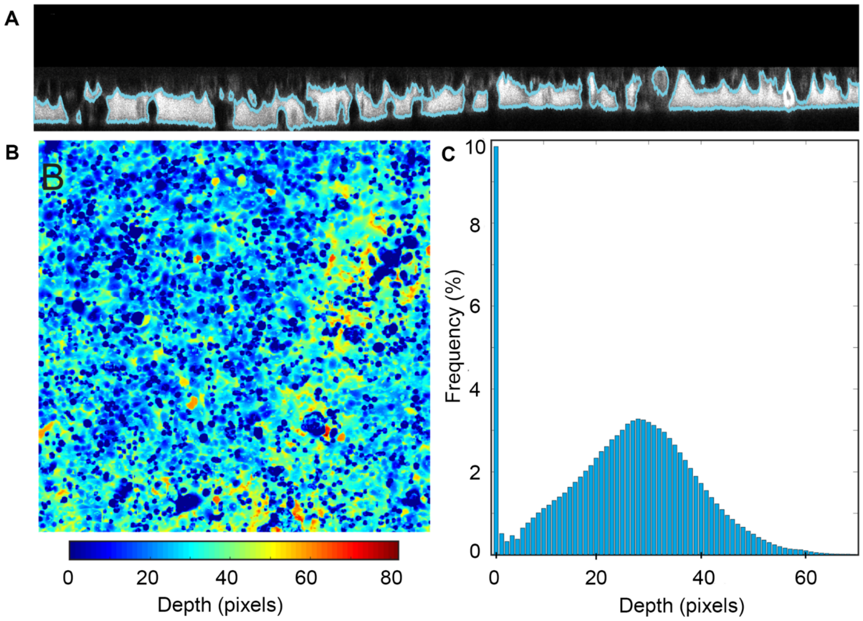


**Supplementary Fig. S8. Method for airway surface liquid (ASL) thickness**. **A)** Sample of an (x, z) cross-section showing the original image and the segmented regions. **B)** Heat map showing thickness across the sample. **C)** Histogram showing the pixel thicknesses across the region in panel B.

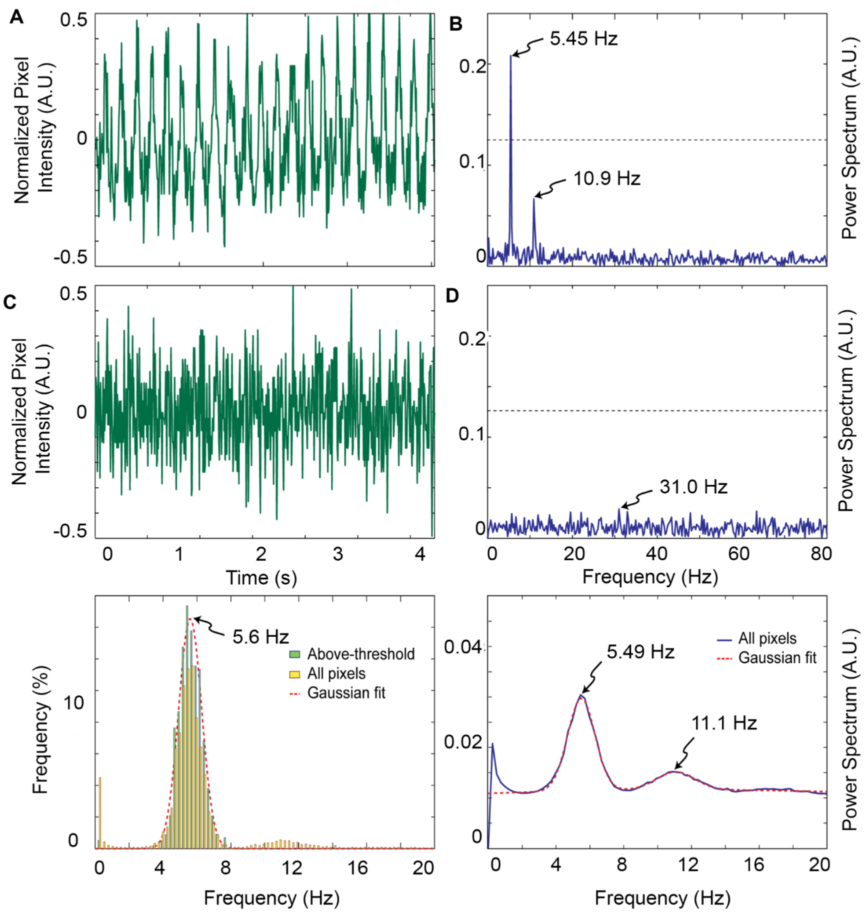


**Supplementary Fig. S9. Method for computing ciliary beating frequency (CBF). A)** Data from two different pixels from a video was normalized so that it ranged from 0 to 1 and then readjusted to have zero mean. **B)** These time courses were used to compute the single-sided spectrum for each pixel, shown for the two pixels in panel A. To consider only pixels with considerable oscillatory behavior (*e.g*. top pixel in panel A, but not bottom), a threshold of 0.125 A.U. was set for the maximum power, shown by the dotted lines. **C)** The frequency where the maximal power was observed for each of the pixels was computed, and this plotted as a histogram. Shown are the histograms for all pixels, and only for those with maximum pixel intensity above the threshold. The latter was fit to a single Gaussian, shown by the dotted line. This was used to determine the average beating frequency for the sample. **D)** As an alternative check, the power spectrum over all pixels was computed, and fit by a Gaussian mixture model (red dotted line).

**Supplemental Video Legends**

**Supplementary Video 1.**

Kliment et al. Suppl Video 1_CTRL pre-CS

Ciliary beat frequency (CBF) in NHBE cells (control adenovirus) before exposure to CS.

**Supplementary Video 2.**

Kliment et al. Suppl Video 2_CTRL post-CS

CBF in NHBE cells (control adenovirus) 30 min after exposure to CS, showing a slowing of CBF.

**Supplementary Video 3.**

Kliment et al. Suppl Video 3_ANT1 pre-CS

CBF in NHBE cells (ANT1-GFP adenovirus) before exposure to CS.

**Supplementary Video 4.**

Kliment et al. Suppl Video 4_ ANT1 post-CS

CBF in NHBE cells (ANT1-GFP adenovirus) 30 min after exposure to CS, showing a slowing of CBF.

**Supplementary Video 5.**

Kliment et al. Suppl Video 5_ ANT2 pre-CS

CBF in NHBE cells (ANT2-GFP adenovirus) before exposure to CS.

**Supplementary Video 6.**

Kliment et al. Suppl Video 6_ ANT2 post-CS

CBF in NHBE cells (ANT2-GFP adenovirus) 30 min after exposure to CS, showing preservation of CBF.

**Supplementary Video 7.**

Kliment et al. Suppl Video 7_Air treated mouse lung

SIM imaging of air-treated mouse lung stained for ANT1 (red, Rabbit anti-human ANT1 antibody, ab102032), tubulin (light blue) and nuclei (dark blue). ANT1 localizes to the plasma membrane surface in a linear pattern on the apical cell surface and to cilia in a striated pattern. ANT also localizes to mitochondria in the cell body.

**Supplementary Video 8.**

Kliment et al. Suppl Video 8_Smoke treated mouse lung

SIM imaging of cigarette smoke-treated mouse lung stained for ANT1 (red, rabbit anti-human ANT1, ab102032), tubulin (light blue) and nuclei (dark blue). ANT1 localizes to the plasma membrane surface in a linear pattern on the apical cell surface and to cilia with loss of patterning. ANT also localizes to mitochondria in the cell body.
